# Habitat filtering but not microbiota origin controls microbiome transplant outcomes in soil

**DOI:** 10.1101/2024.12.12.627922

**Authors:** Senka Causevic, Janko Tackmann, Vladimir Sentchilo, Lukas Malfertheiner, Christian von Mering, Jan Roelof van der Meer

## Abstract

Human activities cause global losses of soil microbiome diversity and functionality. Microbiota transplants offer a potential solution, but the factors influencing transplant success remain unclear. We investigated how microbiota origin affects microbiome mergers, hypothesizing that native strains through niche preference are better adapted to their habitat and will outcompete non-native ones. To test this, we contrasted transplants between soil microcosm-cultured topsoil or lakewater communities with a community of 21 soil bacteria (SynCom). In both cases, SynCom transplant increased resident productivity but permanently shifted compositions, although its abundance dropped from an initial 50-80% to <1% within two months. Both merged and non-merged communities resembled natural soil microbiota in comparisons with over 81,000 soil, sediment and lake compositional data. Our results show that habitat filtering and niche competition, more than microbiota origin, determine transplant outcomes. Despite the limited proliferation of SynCom transplants, their capacity to instill lasting community trajectory changes opens new paths for microbiome engineering.

**TEASER:** Even transiently present microbiota transplants can alter resident microbiome composition through processes governed by habitat filtering.

## INTRODUCTION

Soils encompass staggering microbial diversity and biomass, which sustain a diverse plant and animal life (1, 2). Soil microbiomes (i.e., the ensemble of microorganisms and their activities within the soil habitat (3)) largely contribute to soil fertility by transforming and building soil organic matter, establishing soil aggregate formation through exudates, and facilitating nutrient provision to plants. Collectively, soil microbiomes carry out crucial roles in the major planetary biogeochemical cycles (4, 5). Alarmingly, soil microbiome diversity, soil organic matter and soil structure are threatened world-wide as a result of current agricultural practices, pollution, poor land management, erosion and urbanization (6, 7). Degraded soil structure and microbiome diversity loss will affect future food production (7), land ecosystem stability and climate (8), urging the necessity to reverse the trends of soil degradation. However, soil restoration approaches must be accompanied with fundamental knowledge on the processes underlying assembly of taxa-complex soil microbiomes, and an understanding of the causes leading to mal- or dysfunctional system states.

One of the potential approaches to regenerate degraded soils involves restoration of soil microbiomes (9–11). This can take the form of application of single inoculants with assumed beneficial character, in some case in combination with specific prebiotic substrates (12, 13), but single bacterial or fungal inoculants are unlikely to overcome the multiple factors, regional properties and scales of soil loss (globally one third of all soils are considered degraded) (14,15). Alternatively, defined strain consortia with one or more specific functional guilds could be applied, but there is currently insufficient knowledge to ensure their efficacy. Similar to clinical gut microbiome interventions, one could imagine a complete reset of the impacted soil microbiome followed by habitat recolonization from a community transplant originating from a healthy site. In contrast to the gut, where antibiotics can be applied to diminish the abundance of existing resident bacteria before adding the transplant, resetting a soil microbiome is not achievable at scale. Hence, any soil transplant inoculum would have to compete with the resident microbiota of the impacted soil, but the mechanisms and processes underlying transplant outcomes are poorly understood.

The merger of two distinct microbiomes (for example, a soil transplant with a soil containing a resident community) is known as microbiome coalescence (16). Coalescence has remained surprisingly understudied considering its common occurrence (17). For example, falling leaves, animal droppings, river estuaries or handshakes all cause microbes to disperse from one habitat to another, exerting coalescence (16). In the context of soil, several studies have shown positive effects of transplants on plant disease suppression, drought tolerance or climate tolerance of planted trees (18–20). However, the processes underlying transplant mergers with resident microbiota are unknown, and there is little current theory to predict coalescence outcomes and resulting taxa changes (21). To start building testable hypotheses, it is crucial to use controlled systems with highly reproducible communities, which limit the confounding effects of biotic and abiotic variables present in diverse natural soils. We have recently shown how both defined (e.g., from cultured isolates) and taxa-complex but undefined soil microbiota (using the washed microbial cell fraction from native soil as inoculum) can be grown in microbe-free soil (22). Both inoculum types mature reproducibly on a time scale of weeks to months into compositions with clear soil community signatures (22), making them ideal resources to systematically investigate the process of microbiome coalescence. Here, we focused on delineating the effects of habitat, nutrient niche preference, taxa origin and diversity in establishing community signatures and merger outcomes.

In order to be able to correctly follow individual taxa development in community mergers, we cultured two microbiomes from different origins in soil microcosms, which were transplanted with a defined low-complexity (21 strains) synthetic soil community. Resident and transplant communities were cultured in identical microbe-free soil microcosms for up to a month, as in Ref. (22), during which we characterized their compositional trajectories. To produce soil microbiomes with similar, but distinct taxa compositions we inoculated microcosms with either the washed microbial cell fraction of a top forest soil layer (SoilCom), or the cells recovered from freshwater lake by filtering (LakeCom). We chose freshwater lake microbiota inoculum, since lakes receive soil run-off and hence should still carry taxa similar to soil that could develop into a soil microbiome under habitat selection while having distinct taxa memberships and niche preference (23, 24). The 21 SynCom isolates cover four major bacterial phyla native to the soil and originate from the same site as the SoilCom inoculum, and they reproducibly colonize fresh soil to high density (22). One-month old soil-grown communities were used for transplant experiments, by mixing equivalent soil portions of SynCom and SoilCom or LakeCom into fresh soil microcosms. Importantly, this allows renewed colonization of the fresh soil material and creates nutrient competition between the resident (SoilCom and LakeCom) and the transplant (SynCom) members. To contrast the effect of nutrient availability, we further transplanted fresh SoilCom inoculum into soil-grown SynComs at different stages of maturation (two weeks and three months). Contrary to our expectations, transplanted SynCom members declined in presence of both resident communities, independently of their origin, but its transient presence resulted in lasting compositional changes of both SoilCom and LakeCom. Regardless, a global comparison to over 81,000 soil and lake microbiota compositional data further showed that all resident and transplanted microcosm-grown communities developed typical soil-microbiome signatures, indicating the dominant filtering role of the habitat on the coalescence process.

## RESULTS

### Soil and lake water inocula develop into soil-like communities with distinct productivities in soil microcosms

To produce different resident soil communities to be used in transplant studies, we extracted two natural communities to be used as inoculum for colonization of sterile soil microcosms. The first of these consisted of microbial cells washed from forest topsoil (–5 to –15 cm, named here *SoilCom*), whereas the second was comprised of cells collected from a freshwater lake (0 to –1 m, or *LakeCom*). Since freshwater lakes receive run-off from nearby soils, we hypothesized that LakeCom overlaps to some degree in its taxa composition with SoilCom, and may, therefore, develop into a soil-like microbiome when given the appropriate growth environment. If successful, this would also demonstrate the filtering role of the habitat in directing the development of complex microbiota. Indeed, there was 27.7% taxa overlap at genus level (after removing singletons and non-assigned taxa) between lake and soil inocula, whereas 1% of the detected ASVs in lake inoculum were also found in the soil inoculum.

Both communities were inoculated at the same density in microbe-free soil microcosms (at ca. 10^6^ cells per g soil) and were incubated in parallel for 21 days at ambient temperature in the dark (Fig. 1A). The community sizes increased rapidly during the first three days to more than 100-fold their inoculated densities, indicating active growth (Fig. 1B and 1C). Viable (colony forming units, CFU) and total (flow cytometry) cell counts produced similar trends, although LakeCom CFU counts tended to be lower than the corresponding flow cytometry counts. This may be due to the use of a single nutrient medium for CFU counting, on which multiple taxa may not grow (Fig. 1B and 1C). Both soil and lake-derived communities reached maximum densities between day 3 and 7, after which their sizes slightly decreased and stabilized until day 21. In contrast to our initial expectations, LakeCom community sizes remained 3–10-fold lower than those of the SoilCom (Fig. 1B and 1C, two-way repeated-measures ANOVA on ranked values for the effect of community and incubation time, *P*-values indicated on the figure for the effect of community, Supplementary Data). This suggested that the LakeCom inoculum could not exploit the available nutrient niches in the soil microcosms as well as the SoilCom inoculum.

**Fig. 1.**
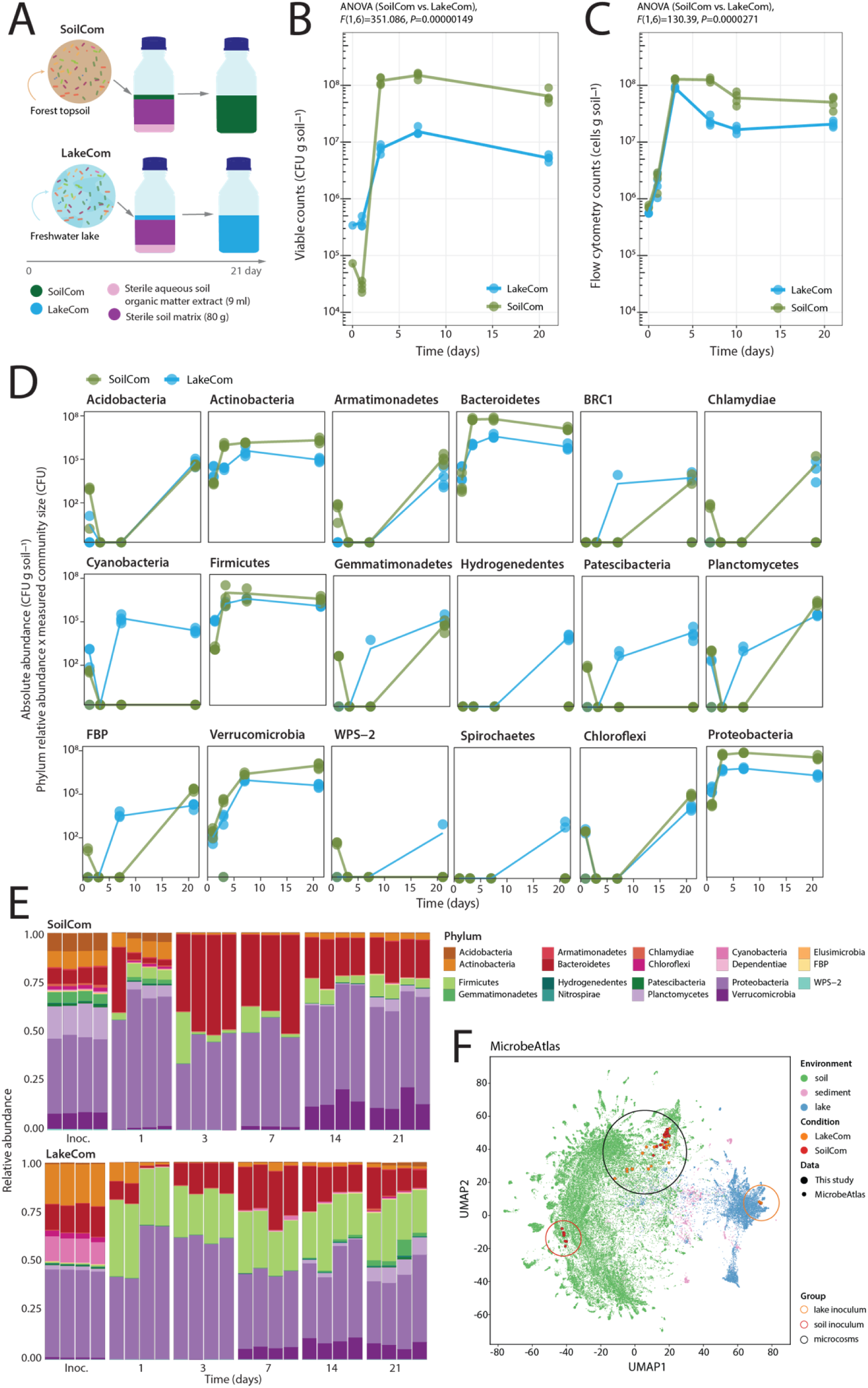
Development of soil and freshwater lake communities in soil microcosms. **(A)** Experimental approach to grow soil (SoilCom) and freshwater lake inocula (LakeCom, both seeded at ca. 10^6^ cells per gram) in soil microcosms. Changes in community sizes over time in the soil microcosms of SoilCom (in green) and LakeCom (in blue), estimated by viable counts (**B,** colony forming unit, CFU, counts per gram of soil) or total community counts by flow cytometry **(C)**. Lines connect the means of the four biological replicates per condition (presented as dots). *P*-values relate to the effect of community type on attained population sizes, as obtained with two-way repeated measures ANOVA on ranked values (Supplementary data). **(D)** Growth of individual phyla in both communities, multiplying the relative proportion of each phylum obtained from gene amplicon sequencing by the total viable community size (CFU), from (B). Lines connect the means from the four biological replicates (dots) per condition (SoilCom in green or LakeCom in blue). **(E)** Phylum-level community compositional changes over time and per biological replicate (relative abundances from sequence data, stacked). Phyla with relative abundances below 0.01 are not shown. **(F)** Uniform Manifold Approximation and Projection (UMAP) for dimension reduction of SoilCom and LakeCom inocula (red and orange circle, respectively), and soil microcosm samples (black circle) on all publicly available soil (green dots), sediment (pink dots) and lake (blue dots) microbiome compositions from the MicrobeAtlas database.

Phyla-proportionalized community growth (CFU counts normalized by V3-V4 16S rRNA gene amplicon sequence relative abundances) indicated that the fast increase during day 1 to 3 is driven mainly by members of four phyla: Proteobacteria, Firmicutes, Bacteroidetes and Actinobacteria (Fig. 1D). The observed differences in SoilCom and LakeCom sizes at those sampling times are also mostly attributed to members of these four phyla (Fig. 1D). Their rapid expansion was accompanied by slower growth of, notably, Verrucomicrobia and other minor phyla, in both inoculum types, and visible at later timepoints. In contrast to the dominant phyla, various minor phyla started from lake inoculum managed to establish better in soil microcosms than the same ones from soil – at least during the incubation time of 21 days. These included Cyanobacteria, Patescibacteria, Gemmatimonadetes, Hydrogenedentes, WPS-2 and Spirochaetes (Fig. 1D). The same trend was observed with phylum abundances calculated from flow cytometry data (Fig. S1A). Proportionally speaking, Firmicutes contributed more than Bacteroidetes to early compositional changes in LakeCom-compared to SoilCom-originated communities (Fig. 1E) and were able to retain these high initial fractions for the whole incubation period.

Despite these differences, both types of soil-grown microbiota showed distinct soil, rather than lake compositional signatures in a comparison to a global background of 81,627 soil, lake and sediment communities from MicrobeAtlas (25) (Fig. 1F). SoilCom and LakeCom compositions converged but clustered separately from their respective inocula, each of which resembled the microbiota compositions typical of their origins (Fig. 1F). Closer inspection of the MicrobeAtlas projection suggested that SoilCom and LakeCom samples fall in an intermediary zone with soil communities characterized by large relative abundances of fast-growing taxa such as Pseudomonas and Pedobacter (Fig. S1B – S1G), which is consistent with growth of these genera in the SoilComs and LakeComs (Fig. S1H and S1I). These results thus showed that both inocula, regardless of their taxa origin (i.e., soil or lake) develop into soil-like communities when grown in microbe-free soil microcosms, showing the strong role of habitat filtering in establishing community signatures. However, their compositional starting differences may have resulted in differing succession trajectories, which in case of LakeCom allowed better proliferation of a variety of minor phyla representatives and in case of SoilCom a higher overall community productivity.

### Common freshwater lake and soil genera display different productivities depending on the community development trajectory

To better understand which taxa and developmental stages were responsible for the observed differences in growth of SoilCom and LakeCom, we looked deeper at their compositional differences. Communities from both origins displayed an initial decline of Shannon diversity (as a result of fast-growing blooming populations combined with limited sample sequencing depth), and a subsequent recovery (Fig. 2A). Despite starting at a lower apparent diversity, LakeCom microbiota in soil microcosms developed higher detectable taxa diversity than SoilCom after 21 days of incubation (Fig. 2A, two-way repeated measures ANOVA for community and time effect, *F*(5,25)= 38.077, *P*-value=6.59×10^-11^, *post hoc t*-test at day 21, *P*=0.00026, Fig. S2A). Both community types maintained significantly different compositions throughout the incubation (Fig. 2B, ASV level; PERMANOVA, *P-* value=0.001 for both community type and time effect, Supplementary Data). However, this difference is less visible (while still statistically significant) at higher taxonomy levels (e.g., Family, Fig. S2B), likely due to habitat filtering for similar functional groups. Indeed, there was an increase of shared ASVs among LakeCom and SoilCom from 1% at the inoculum stage (see above) to 12% after 21 days incubation in the soil microcosms.

**Fig. 2.**
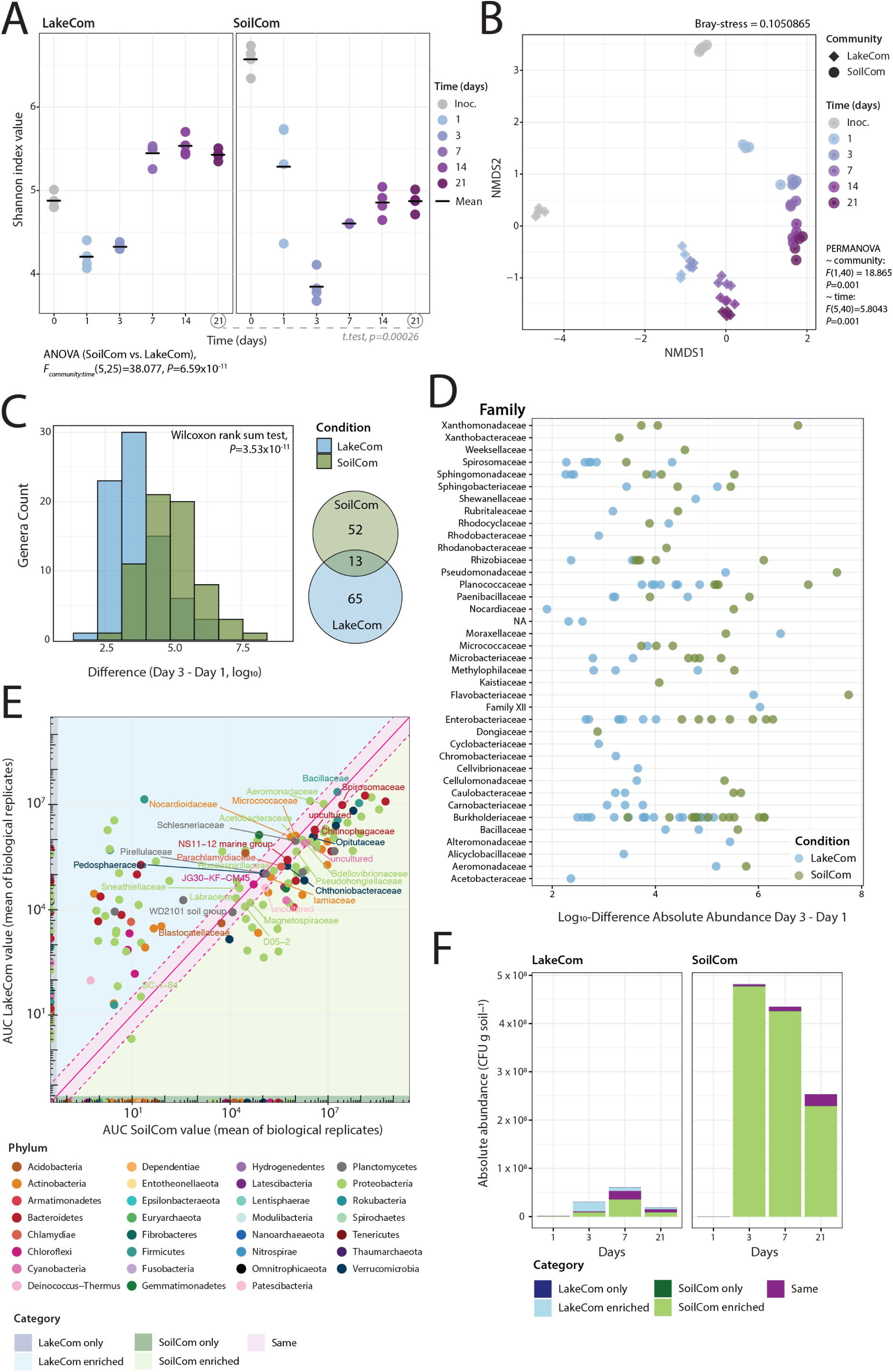
Differences in diversity and productivity of soil and freshwater communities in soil microcosms. **(A)** Shannon diversity changes of SoilCom and LakeCom communities over time (4 biological replicates per community). Black bars indicate the mean of the four replicates, with circles indicating individual values. Time progression is illustrated with darkening colours as shown in the legend. The effect of time and community on index values were tested with two-way repeated measures ANOVA. Dashed line shows *P*-value of *post hoc t*-test comparing the last time point of growth. **(B)** Non-metric multidimensional scaling (NMDS) ordination of LakeCom (diamond shapes) and SoilCom (circles) samples using Bray-Curtis distances on ASV-level compositions. Darkening colours depict time progression as in (A). The effect of community and time on pairwise distances was evaluated with PERMANOVA (Supplementary Data). **(C)** Differences in the number of early-colonizing fast-growing genera (i.e., those that increased their absolute abundance from day 1 to day 3) and the extent of their increase (log_10_ transformed) in LakeCom (in blue) and SoilCom (in green; shown as histograms). The Wilcoxon rank sum test compares both distributions. The Venn diagram on the right shows the number of unique and overlapping early colonizers between SoilCom and LakeCom. **(D)** Log_10_-absolute abundance differences between day 3 and day 1 for each of the genera from (C) (circles), grouped by their family-affiliation, and coloured by SoilCom (green) or LakeCom origin (blue). **(E)** Comparison of family growth (summed for all detected ASV per family; not only early colonizers), between LakeCom and SoilCom. Circles show paired means of the area under the curve family productivity (AUC, using absolute abundances from CFU data) for each of the communities (SoilCom on the abscissa, LakeCom on the ordinate). Circle colours show phyla affiliations. Coloured areas in the graph highlight the defined AUC-comparison categories, according to the colour legend below. Families exclusive for LakeCom are highlighted in dark blue; those exclusive to SoilCom in dark green; families with similar AUCs in the magenta area; light blue for families enriched in LakeCom, and light green the families enriched in SoilCom. **(F)** Summed absolute abundances of families in each category from (E) (same colours, darker shades) at individual time points in LakeCom and SoilCom.

To detect taxonomic differences among early colonizers that may have influenced the subsequent growth trajectories of the complete communities, we selected genera in both resident communities with an increase in absolute abundance between day 3 and day 1 (Fig. 1B, based on CFU counts). This resulted in identification of 65 unique genera in LakeCom and 52 in SoilCom communities, whereas 13 fast-growing early colonizing genera were found in common (Fig. 2C). Despite the higher number of early colonizing genera in LakeCom, their extent of growth was on average lower than those from SoilCom (Fig. 2C, Wilcoxon rank sum test, *P-*value=3.53×10^-11^). Grouped by families, and with few exceptions, most early colonizing genera from SoilCom were more productive in the soil microcosms than those from LakeCom (Fig. 2D). Very few genera belonged to families exclusive for either the SoilCom (e.g. Xanthomonadaceae, Xanthobacteraceae), or the LakeCom origins (e.g. Rhodobacteraceae, Shewanellaceae, Fig. 2D). The higher productivity by SoilCom early colonizers was less evident (and not statistically significant) with absolute abundances based on flow cytometry counting (Fig. S2C, S2D), due to measured high LakeCom productivity at day 3.

A comparison of both systems based on the summed productivities of all families across all time points, defined as the area under their growth curve (AUC, absolute abundance from CFU data; as in Fig. 1D), indicated five distinct taxa subsets. The first subset included families with similar AUC values in both SoilCom and LakeCom, for instance, the Pedospheraceae, Pirellulaceae, Bacillaceae, or Aeromonadaceae (Fig. 2E, the diagonal area). Two other subsets encompassed families exclusive to either LakeCom (*LakeCom-only*) or SoilCom (*SoilCom-only*, Fig. 2E). Finally, two subsets covered common families enriched in LakeCom microcosms but poorer in SoilCom (*LakeCom-enriched*), or vice-versa (*SoilCom-enriched*, Fig. 2E).

Surprisingly, the exclusive subsets of either SoilCom and LakeCom contributed only marginally to the overall community sizes (Fig. 2F; LakeCom-only, SoilCom-only). In contrast, among the shared families, it was mostly the difference in productivity of the SoilCom-enriched subset that resulted in the much larger community size of SoilCom than LakeCom (Fig. 2F). Calculating AUC-values at genus or family level but from flow cytometry-inferred absolute abundances produced the same trends (Fig. S2E, 2F), indicating that it is less the absence of specific families in LakeCom that limited its community growth in soil microcosms, but rather their lower productivity. Excluding the possibility of a disproportionate importance of the SoilCom- and LakeCom-only fractions (e.g., key taxa missing from lake, but present in SoilCom-only fraction and crucially important for some nutrient transformation), LakeCom lower productivity can thus be attributed to taxa and metabolic differences at genus (or species) levels.

### Merging soil-native and non-native communities with a synthetic soil community changes their developmental trajectories and productivity

Given that both LakeCom and SoilCom had colonized the soil microcosms to their maximum potential, and had developed into reproducibly distinct soil microbiomes, we proceeded with the coalescence stage. Since LakeCom was less productive than the SoilCom in the soil microcosms (Fig. 1B, 2A), we expected that the presence of the culturable soil taxa from the SynCom would redirect the developmental trajectory of LakeCom to a higher productivity state. In contrast, since SynCom members originate from the same source material as SoilCom, we expected them to have no effect on merged SynCom+SoilCom growth. To test this, we mixed a one-tenth weight fraction of either of the one-month old soil-grown communities with the same quantity of soil from microcosms colonized for 1 month by the defined SynCom and with eight times that weight of freshly sterilized soil microcosm material, to allow the merged communities to recolonize the soil (Fig. 3A). As further comparison for the developmental trajectories, we prepared control soil microcosms with similarly diluted SoilCom or LakeCom but without SynCom addition, and with diluted SynCom by itself.

**Fig. 3.**
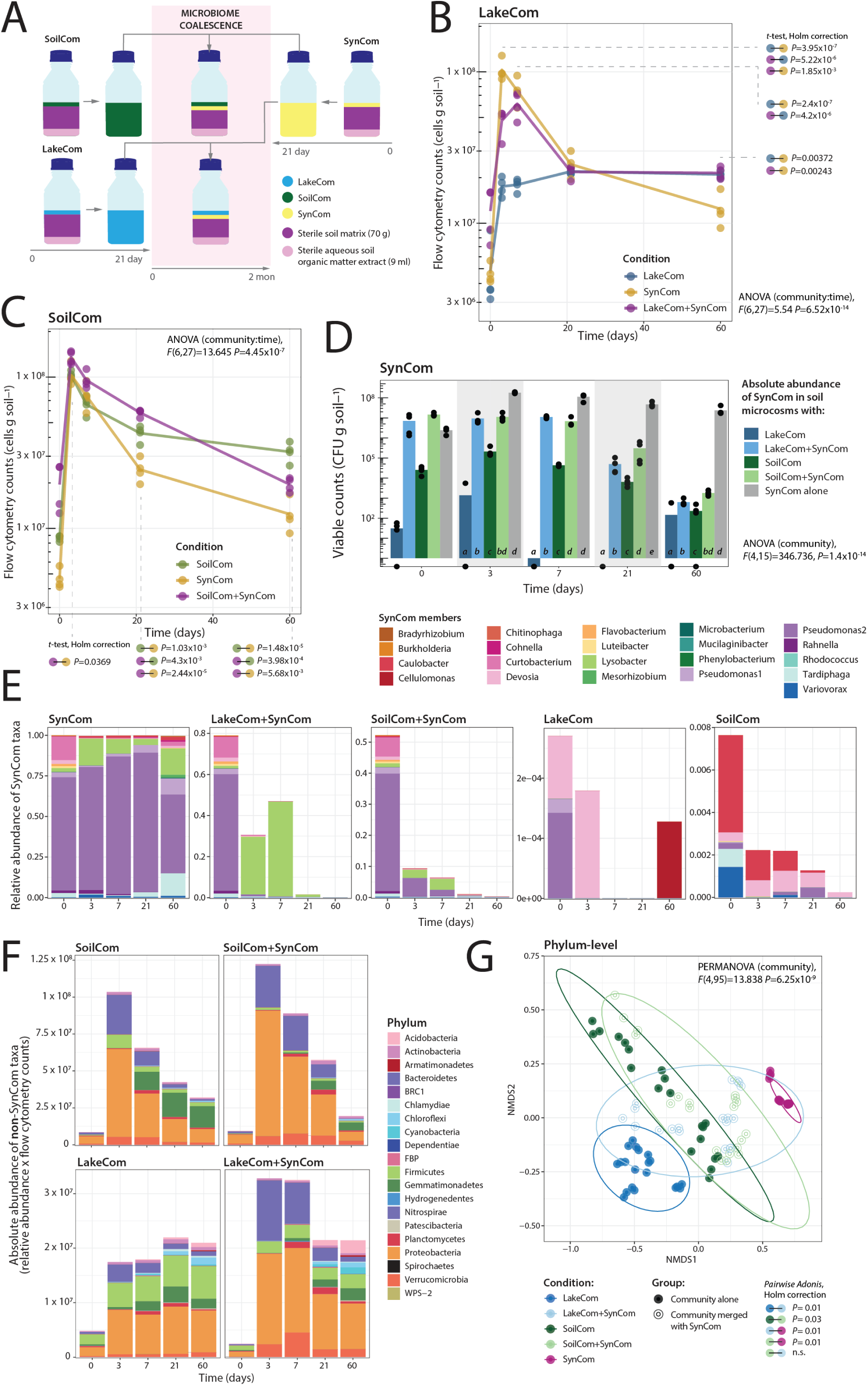
Coalescence with SynCom changes LakeCom and SoilCom growth and composition. **(A)** Design of the coalescence set-up. Stabilised SoilCom or LakeCom are mixed with stabilised SynCom in a 1:1 soil-weight ratio and supplemented with 8-fold sterile soil with fresh supply of soil organic extract in new microcosms. Controls consist of individual diluted communities in new microcosms. Flow cytometry counts of merged and individual community sizes, for LakeCom **(B)** and SoilCom mergers **(C)**. Individual biological replicate values appear as circles, with lines connecting the means. Blue colours, LakeCom alone, Green colours, SoilCom alone; magenta, merger with SynCom; yellow, SynCom alone. Two-way repeated measures ANOVA on ranked values tests for community and time effects. *Post hoc t*-tests show paired comparisons at specific time points (Holm’s correction, *P*-values only shown if below 0.05). Connected circles and colours on the right side of the plot indicate the comparison pair. **(D)** SynCom summed community sizes across all conditions. Community size represented as total viable counts per gram, multiplied and summed from the relative SynCom ASV abundances and the total community CFU counts in each sample and condition. Black circles show biological replicate values and bars show their means (n = 4 biological replicates). Bar colours represent community type as per the panel legend. Letters show significance groups, for each time point shown separately (except t=0), obtained with pairwise *t*-tests (Holm’s correction) following a significant *P*-value of repeated measures ANOVA on ranked values. **(E)** Mean relative abundances of the SynCom members (within the summed fraction displayed in D) as proportion of the total community composition. Colours on the stacked bar plot indicate SynCom taxonomic membership as per the legend shown above the panel. **(F)** Absolute phyla abundances of the non-SynCom fraction per community type over time (calculated as the remaining proportion from (E), multiplied by the total community size as measured with flow cytometry. Phyla colours indicated on the right of the panel. **(G)** Non-metric multidimensional scaling (NMDS) ordination of paired Bray-Curtis distances of phylum-level compositions of all merged and non-merged communities. Circles group all time points and replicates per community type, as indicated by the different colours below the panel. Ellipses connect multivariate t-distributions of the community types (same colour code). PERMANOVA tests significance of community type effect, with individual paired *post-hoc* comparisons shown below the panel (connected circles and colours indicate paired comparison groups; Supplementary Data).

As in the first incubation phase, growth was observed for all communities during the first three days after mixing the soil into fresh soil microcosms (Fig. 3B, 3C, S3A, S3B). The size of the merged SynCom+LakeCom community indeed surpassed that of the non-merged LakeCom controls at days 3 and 7, but this effect levelled off after day 20 (Fig. 3B). SynCom mergers with SoilCom also resulted in a slightly higher community sizes than SoilCom by itself, but the effect was less prominent than for LakeCom mergers (Fig. 3C). Eventually, the sizes of SoilCom+SynCom mergers and of the SynCom community by itself decreased to below those of SoilCom alone (Fig. 3C).

Although the sizes of both SynCom-merged communities were higher than of their non-merged controls at day 3 and 7 (Fig. 3B, 3C), the SynCom members in those communities showed no net growth, whereas the SynCom community by itself did (light blue and light green bars compared to grey bar, Fig. 3D). Rather, the SynCom proportion decreased at a first-order rate relative to the total community size, from 52–79% at day 0 to 9–30% at day 3, and down to 1% at day 20 and below (Fig. 3E, height of stacked bars). Detectable SynCom-specific ASVs naturally present within the unmerged SoilCom (since they originate from the same soil sample) increased their population sizes up to day 3, beyond which they declined similarly as in SynCom mergers (Fig. 3D). The relative abundances of SynCom members differed between both merged communities, as well as between non-merged SoilCom and SynCom (Fig. 3E). Notably, within the LakeCom merger, populations of the SynCom-member Lysobacter bloomed, whereas this was much less pronounced in the SoilCom merger and within SynCom by itself, where two pseudomonads made up the dominant populations (Fig. 3E). SynCom absolute abundances at day 60 remained slightly higher in the merged than in the non-merged microcosms at ca. 1600 and 500 CFU per gram for soil and lake merger condition, respectively (Fig. 3D, results of *post hoc t*-test after a significant two-way repeated measures ANOVA on ranked values; see Supplementary Data), but with different compositions among the systems (Fig. 3E).

We additionally studied the mergers with fast expectation-maximization microbial source tracking (FEAST) that quantifies the presence of the source communities at each time point. This analysis showed that both SynCom and either LakeCom or SoilCom can be detected in the merger up to day 7, after which the fractions of sourced SynCom in the mergers decline to below detection (Fig. S3C and S3D). Interestingly, an ‘Unknown’ fraction not attributable to either source appeared over time, which is reflecting the compositional shift in the mergers following the SynCom transplant (Fig. S3C and S3D). Comparison of the merged versus non-merged community compositions (absolute taxa abundances based on relative ASV abundances and normalized between the samples by flow cytometry counts, excluding SynCom-members), indicated a notable decrease in the relative abundance of Firmicutes within LakeCom as a result of SynCom merging, and a consequent increase in Proteobacteria, Bacteroidetes and Verrucomicrobia (Fig. 3F). SoilCom by itself differed from the merged SoilCom+SynCom by the increased proportion of Gemmatimonadates (Fig. 3F). Compositional trajectories in non-metric multidimensional scaling ordination were significantly different between merged and non-merged communities on the same taxonomy level (Fig. 3G, PERMANOVA analysis, for LakeCom: *P*_LAKE_-value=0.01; for SoilCom, *P*_SOIL_-value=0.03, Supplementary Data, Fig. S3E). However, merged communities were not significantly different between each other (Fig. 3G, phyla-level comparison, SynCom reads kept in the data set, SoilCom+SynCom vs. LakeCom+SynCom). Collectively, these results indicated that SynCom transplants temporarily increased global community productivity and changed the developmental trajectories of LakeCom and SoilCom, even though eventually very little remained of the inoculated SynCom members.

### SynCom transplants impose distinct changes to merged partners compositions, but resulting communities remain soil-like

To describe how community development trajectories changed at higher taxonomic resolution in the merged compared to non-merged communities, we analysed ASV-level compositional differences over time, excluding the SynCom-ASV reads. Non-metric multidimensional scaling ordination separated (merged and non-merged) LakeComs from SoilComs, with 47% of variation explained by community type and 22% by time (PERMANOVA, Fig. 4A). The effects of merging on compositional variation for both LakeCom and SoilCom were clearest in the early time points (0 to day 7), but were statistically significantly different across the complete time series (Fig. 4A, pairwise *adonis*, *P-*value=0.006 for each of the merged community vs. its non-merged control; Supplementary Data). Merger effects are also detected in the transient changes of diversity measures of LakeCom and SoilCom (Fig. S4A). Even though SynCom members themselves declined in the coalesced LakeCom and SoilCom (Fig. 3D), their transient presence and/or activity thus caused a lasting effect on community compositions (Fig. 4A).

**Fig. 4.**
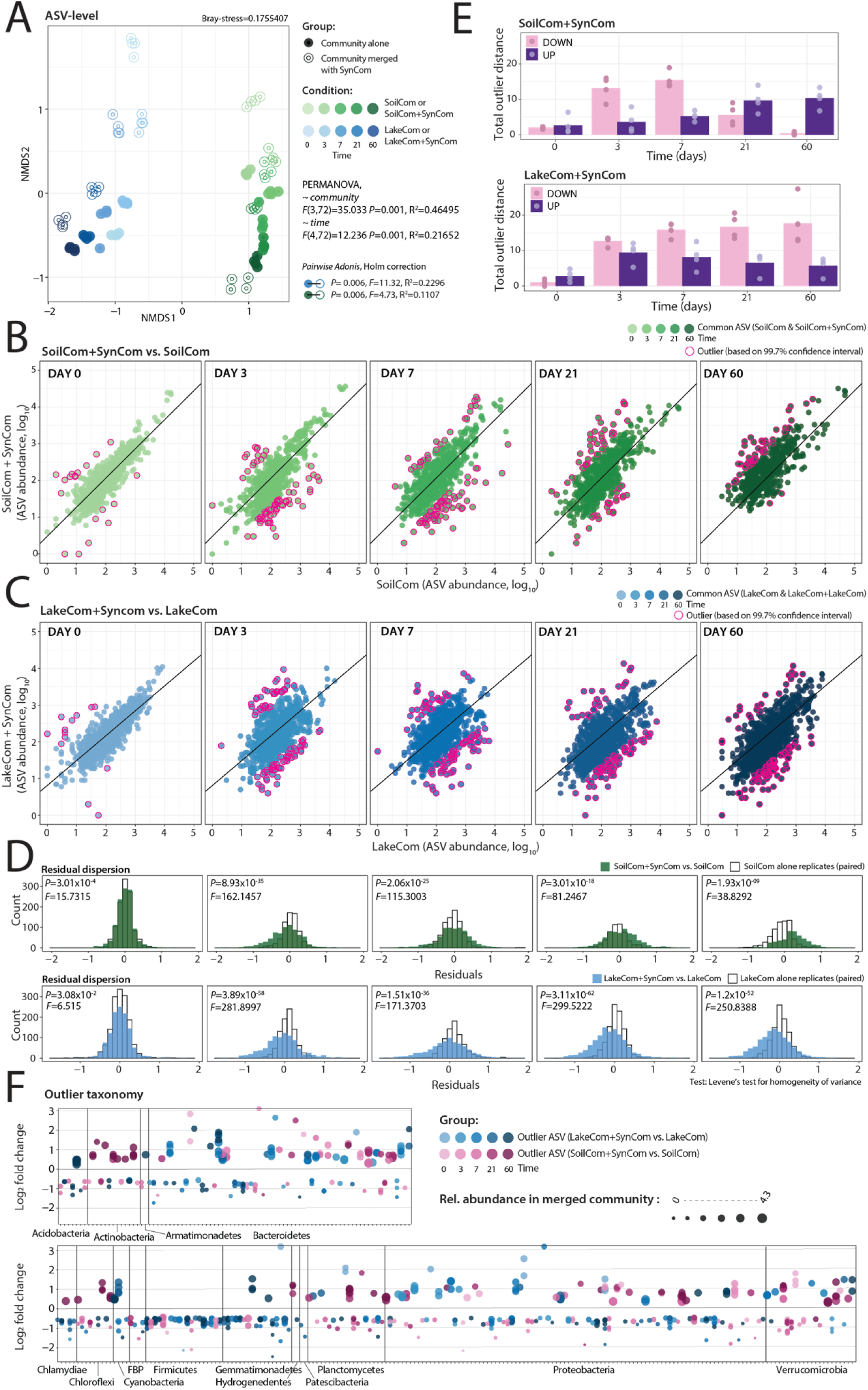
Effects of SynCom merger on the ASV-level composition of LakeCom and SoilCom. **(A)** NMDS ordination of Bray-Curtis distances from pairwise ASV-level sample comparisons among merged and non-merged communities. Circles indicate individual replicate communities, with colour darkness representing time progression (colour legend on the right of the panel). Filled circles, non-merged SoilCom (green) or LakeCom (blue), open circles for merged communities. PERMANOVA tests effects of community type and incubation time. Connected circles on the right side of the plot show pairwise *post hoc* comparison between both merged communities (open circle) and their non-merged controls (filled circle; blue, LakeCom; green, SoilCom; Supplementary Data). Taxa displacement analysis for SynCom merger with SoilCom **(B)** and LakeCom **(C)**. Circles show paired log_10_-transformed ASV abundances in SynCom-merged (but SynCom reads removed) vs. non-merged controls (n=4 biological replicates, paired randomly). ASV abundances paired from equalized datasets, all randomly subsampled to 100,000 reads. Individual plots show outlier ASVs (magenta circles) at each time point, defined as having a residual distance to the t=0 regression line (black line in plots) above the 99.7% confidence interval (derived from the residual dispersion of the t=0 paired data). **(D)** Histograms show residual distribution over time for the merged vs non-merged community comparison (dark colours), and the corresponding non-merged community by itself (random replicate pairs; white colour). *P-* and *F-* values indicate distribution comparisons using Levene’s test for homogeneity of variance. **(E)** Summed residual distances of the outliers shown in (B) and (C), per merger condition (SoilCom+SynCom or LakeCom+SynCom) and direction (‘UP’ for enriched – in dark hue; or ‘DOWN’ for depleted outliers in merged communities, light hue). Circles indicate individual summed replicate values, with bars shoing their means (n = 4 biological replicates). **(F)** Phyla attribution of ASV outliers detected in SynCom-merged LakeCom (hues of blue, darker are later time points) or SoilCom (hues of red). Circles represent the mean log_2_-fold change of the respective outlier ASV abundance in the merged vs. its non-merged community control, with circle size corresponding to its relative abundance in the merged community background (as per legend on the panel top).

To assess the level of disturbance caused by the mergers, we paired common ASV abundances per time point between merged and non-merged communities and quantified the residual variation (SynCom ASVs excluded, see Materials and Methods, Fig. 4B, 4C). Compared to a *null*-model, both SoilCom- and LakeCom-coalesced communities showed considerable taxa displacement over time, even two months after the merger with SynCom (Fig. 4B, 4C). As a threshold for definition of displacement, we used a 99.7%-confidence interval (i.e., ± three times the standard error), calculated from the residual variation among paired ASVs in merged vs. non-merged controls at t=0 (since we assume there is no influence of a merger yet). We verified this further by comparing paired ASV data dispersion among non-merged community replicates alone, which was smaller than between merged-non-merged pairs (Fig. 4D, Levene’s test for homogeneity of variance, *P*-values indicated on the figure, Fig. S4B). Summed over all outliers (i.e., paired ASVs beyond the displacement confidence interval), coalesced SoilCom+SynComs showed an initial increase of depleted ASVs up to day 7, followed by a decrease (Fig. 4E, light colours). In contrast, the sum of specifically enriched ASV outliers increased slowly over incubation time in the SoilCom+SynCom mergers (Fig. 4E, dark colors). LakeCom mergers with SynCom showed the opposite trend, with an increase in depleted ASV outlier distances over time, and a temporal increase and then decrease of enriched ASV outliers (Fig. 4E). Overall, the outlier distance sums (summed distances of both enriched and depleted outlier ASVs to the *null* model) were higher for the coalesced LakeComs than for the SoilComs (Fig. 4E, S4C), and higher than the summed random outlier distances within non-merged LakeCom or SoilCom (Fig. S4C and S4D). This suggested, therefore, a higher degree of disturbance exerted by SynCom on the LakeCom resident community.

The individual outlier ASVs defined from the non-merged vs. merged community comparisons overlapped very little between SoilCom and LakeCom mergers (Fig. 4F). For example, at phylum level, Firmicutes-ASVs were exclusively, reproducibly depleted in both LakeComs and SoilComs upon merging with SynCom, without a single Firmicutes-ASV being enriched. Also, Gemmatimonadetes-ASVs were mostly depleted upon SynCom merging (Fig. 4F). In contrast, at all time points, both enriched and depleted ASV outliers were detected among the Proteobacteria, Bacteroidetes, Planctomycetes, and Verrucomicrobia phyla. Some phyla showed pronounced late responses, for example, the enrichment of ASV outliers in the Actinobacteria and Chloroflexi phyla in SoilCom, and in the Cyanobacteria phylum in LakeCom communities (Fig. 4F).

Finally, we placed the merged and non-merged community compositions in a uniform manifold approximation and projection (UMAP) for dimension reduction, processed similarly at 97% OTU threshold definition with the 81,627 publicly available characterized natural soil and lake communities in MicrobeAtlas. Interestingly and despite the detected taxa displacements at ASV level, both SynCom mergers and LakeCom and SoilCom by themselves located very closely together and showed strong similarity to other soil microbiomes (Fig. 5A). The soil signature drifted from the intermediary zone of the projection in the first incubation phase (Fig. S1B), to a more central position within the soil cluster of MicrobeAtlas in the second and merged communities (Fig. 5B). The central position is marked by higher diversity natural soil microbiomes and suggests maturation of the microcosm communities (Fig. 5C, 2A, S4A).

**Fig. 5.**
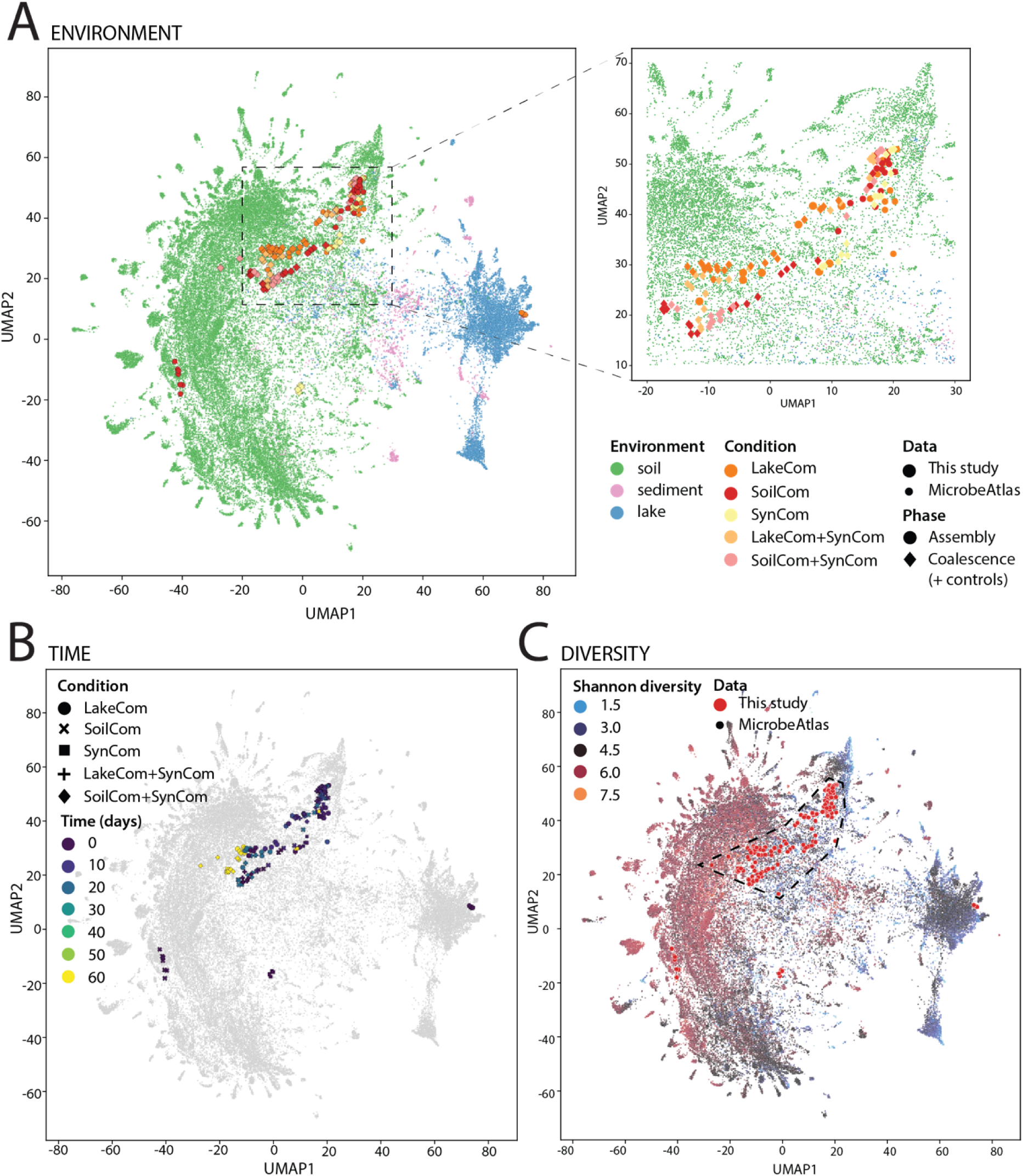
Merged and non-merged LakeComs and SoilComs cluster with natural soil microbiomes from MicrobeAtlas. **(A)** UMAP clustering of all experimental soil microcosm samples (pre- and post-coalescence) and available soil (in green), lake sediment (light pink) and lake water microbiome (blue) samples from the MicrobeAtlas database. Colours and shapes specify different environments and sample origins as per the figure legend. The subplot on the right is an expansion of the main UMAP region with shapes relating to our samples enlarged manually for clarity (indicated with the dashed lines and rectangle). **(B)** As in (A) but highlighting the time progression of the soil microcosm samples. **(C)** As in (A) but showing Shannon diversity levels. The polygon with the dotted black line was added manually for clarity and includes all experimental samples except for the inocula of SoilCom, SynCom and LakeCom.

Taken together, these results thus indicated that the SynCom merger caused pronounced and lasting taxa displacement (on ASV level) in the developmental trajectories of both SoilCom and LakeCom. However, at phylum (Fig. 3G) and 97% OTU comparison within MicrobeAtlas, both non-merged and merged communities, despite their inoculum origin, resembled each other and natural soil microbiomes. We conclude from this that the soil habitat poses the stronger selection on the community development trajectory than the origin of the inoculum.

### SynCom age in soil microcosms determines its colonisation resistance to SoilCom

To identify the potential mechanisms underlying the unexpected and uniform decline of SynCom in mergers (Fig. 3D), we investigated the outcomes of reverse transplants, focusing on SynCom and SoilCom. In this case, we took SynComs at different stage of aging in their soil microcosms, which we expected would influence the availability of nutrients in the systems. Despite its lower taxa diversity (21 members), SynComs grow to similar size in the soil microcosms as the more diverse SoilCom (ca. 10^8^ cells per gram), suggesting that SynCom members occupy all the major nutritional niches. If this was true and nutrient competition was the reason for SynCom decline, we expected soil-grown SynComs at different timepoints of incubation to show varying resistance to invasion with highly taxa-diverse SoilCom. To test this experimentally, we grew SynCom in soil microcosms for either two weeks (“*Syn2w*”) or three months (“*Syn3m*”), after which they were mixed with SoilCom and incubated for one month. Here we used freshly isolated inoculum from the original topsoil at a density of 10^5^ cells per gram, which amounts to ca. 0.1% of the measured SynCom community size. To not include any further nutrients and give SoilCom transplant an advantage, the SoilCom inoculum was applied as cell suspension without any soil or soil nutrient solution (Fig. 6A). As controls, we continued incubation of non-merged SynComs (Syn2w and Syn3m), and we followed SoilCom development from the same washed inoculum in sterile microcosms with only water added instead of soil extract (Fig. 6A).

**Figure 6.**
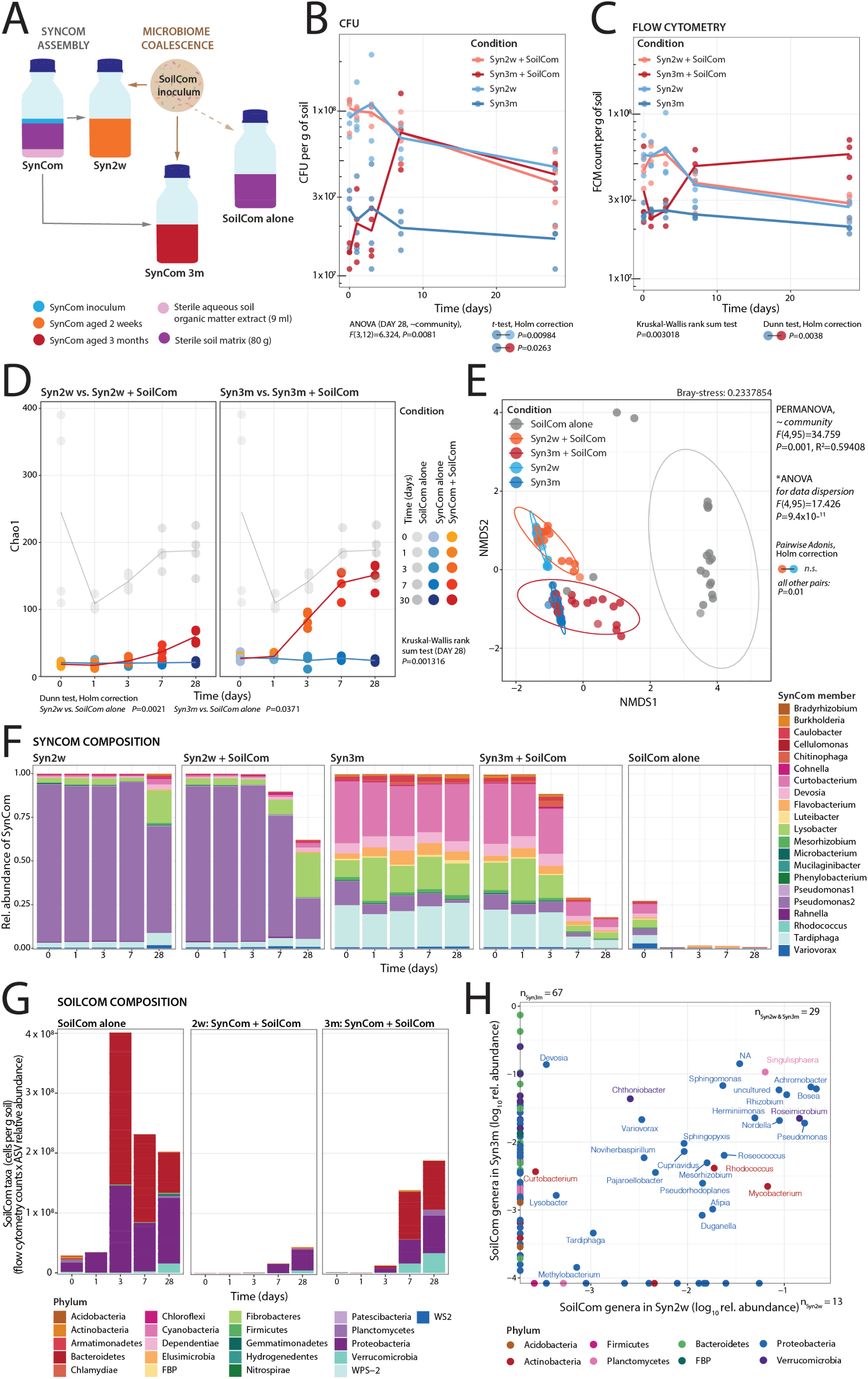
SynCom aging decreases colonisation resistance to SoilCom coalescence. **(A)** Design of the SynCom colonisation resistance experiment. Two weeks (Syn2w) and three months (Syn3m)-aged SynComs in soil microcosms are inoculated or not with fresh SoilCom inoculum (washed cell suspension only) to a density of 10^5^ cells per g and incubated for one month (n = 4 biological replicates). SoilCom alone is the same inoculum inoculated in sterile soil microcosms, but amended with water only. Community sizes over time in merged or control communities, quantified by **(B)** replicate CFU counts per gram of soil, and **(C)** flow cytometry. Circles represent individual replicate values with lines connecting their means (n = 4 replicates). Circle colours represent Syn2w (in light blue), Syn2w+SoilCom (orange), Syn3m (dark blue) and Syn3m+SoilCom (red). Differences between conditions were investigated using one-way ANOVA and *post-hoc t-*tests (Holm’s correction) on the last time point CFU measurement (B), or by Kruskal-Wallis rank sum and *post hoc* Dunn tests (with Holm’s correction) for flow cytometry (C). Only significant *P*-values (<0.05) are indicated. **(D)** Diversity changes (Chao1 values) over time in merged and non-merged control communities. Circles indicate individual replicate values per sample condition (according to colour legend on the side) and time (note that x-axis is discretised), with lines connecting the means. Statistical testing as in (B). **(E)** Relative abundances of SynCom members (coloured and stacked by genus level according to legend next to the panel) across tested communities and time. Stacked bars show the means of relative abundances of four replicates. **(F)** Non-metric multidimensional scaling (NMDS) ordination based on paired ASV-level Bray-Curtis distances. Circles represent individual replicate and time point values, coloured by condition (as per the colour scale within the panel). Ellipses show multivariate t-distribution connection of conditions (same colour code). Differences between communities were tested by PERMANOVA, and ANOVA was used to verify data dispersion. *P*-values of pairwise *post hoc* adonis tests are indicated next to the panel using connected circles to indicate the comparison (Supplementary Data). **(G)** Phyla composition attribution of SoilCom inoculum taxa (as cells per g soil, derived from flow cytometry counts multiplied by ASV relative abundances) in the community setups over time. Stacked bars show means of calculated abundances from 4 biological replicates. Phyla colour code shown below the panel. **(H)** Common and unique SoilCom genera among Syn2w and Syn3m-merged communities at the last time point (day 28; as log_10_-transformed relative abundances). Circles are the means from n=4 replicates, with colors indicating phylum-level affiliation as per the legend below the panel. n, number of SoilCom genera detected exclusively in Syn2w (n_Syn2w_), or Syn3m (n_Syn3m_), or common to both (n_Syn2w&Syn3m_).

The community sizes of both Syn2w alone and mixed with SoilCom inoculum showed a similar, slow decline over one month, both counted by CFUs and by flow cytometry (Fig. 6B and 6C). In contrast, both CFU and flow cytometry counts of the more aged Syn3m communities mixed with SoilCom inoculum increased by two- to fivefold in all replicates, in comparison to the Syn3m control (Fig. 6B and 6C; one-way ANOVA for CFU community effect on last time point values *P*-value=0.0081, *post hoc t*-test Syn3m vs. Syn3m+SoilCom, *P*-value=0.0263, after Holm’s correction. Kruskal-Wallis rank sum test for flow cytometry counts, *P*-value=0.003018 and *post hoc* Dunn test, *P*-value=0.0038 after Holm‘s correction). SoilCom by itself reached a total size of 10^8^ cells per gram of soil, as expected from exploiting inherent soil nutrient sources and thus demonstrating its overall viability (Fig. S5A). These results thus indicated that SoilCom members managed to grow and had found nutritional niches in the Syn3m soil microcosms, but less so in Syn2w.

As expected from the observed merged community growth, a variety of SoilCom taxa established in the Syn3m+SoilCom mergers, which reached Chao1 diversity levels similar to that of SoilCom by itself after one month of incubation (Fig. 6D, see Fig. S5B for a comparison of Shannon index values). However, despite showing no net growth, also merged Syn2w communities increased in terms of Chao1 compositional richness, suggesting at least some SoilCom taxa to establish (Fig. 6D). Correspondingly, the total relative abundance of SynCom membership in Syn3m decreased from 100% to ca. 18% one month after merging with SoilCom inoculum, while that of Syn2w was reduced to ca. 62% (Fig. 6E height of bars). This showed that both tested SynComs were not resistant to invasion from taxa originating from the SoilCom inoculum. Given that Syn3m is more aged than Syn2w, this makes it unlikely that invading SoilCom members after 3 months can use nutrients that somehow were not accessible at 2 weeks of SynCom growth. Rather, it suggests that nutrients were liberated as a consequence of 3 months of SynCom maturation, for example, in form of dead cells (note that the viable SynCom community size after 3 months is one-third of that after 2 weeks, Fig. 2B).

Finally, we examined whether compositional changes could help to explain the higher invasion by SoilCom taxa in Syn3m backgrounds. The compositions of Syn2w and Syn3m differed at the start of coalescence with SoilCom inoculum (Fig. 6B, 6C, 6E and 6F), with higher taxa richness and evenness in Syn3m than Syn2w (Fig. S5B) and notable decrease of Rahnella populations (Fig. 6E). Compositional changes upon SoilCom mergers were more dramatic for Syn3m-than for Syn2w-backgrounds (Fig. 6F, NMDS projections of pairwise ASV-level Bray-Curtis distances; *P*-value=0.01, *post hoc* pairwise adonis after PERMANOVA, Supplementary Data). Invading SoilCom taxa in Syn2w backgrounds consisted primarily of representatives of Proteobacteria, and of Bacteroidetes, Proteobacteria, Planctomycetes and Verrucomicrobia in Syn3m (Fig. 6G). Since there are no Planctomycetes and Verrucomicrobia representatives in the SynCom, their growth could be due to unexploited niches in the soil, although even then, one would have expected these niches to be equally present after 2 weeks and 3 months. At genus level, we identified 29 genera in common between the SoilCom expansion in Syn2w and Syn3m backgrounds, 67 genera unique for Syn3m and 13 for Syn2w (Fig. 6H, Supplementary Data).

In conclusion, these results indicated that resistance of SynCom to invading SoilCom was inversely dependent on its age. Since there is no net SynCom community growth between 2 weeks and 3 months, it is unlikely that the increased SoilCom growth in Syn3m background is a result of non-exploited nutritional niches in the soil microcosms. Rather, the inverse colonization resistance correlated to the lower proportion of live cells in Syn3m. The better growth by SoilCom-taxa in the Syn3m-aged community could therefore be a combination of reutilization of carbon and nutrients from dead SynCom cells, exploitation of non-utilized nutritional niches (as e.g., by the Verrucomicrobia) or lack of some active defence or exclusion mechanism that relates to the proportion of viable cells in the residing community.

## DISCUSSION

To counteract perceived microbiome dysbiosis, intervention strategies using transplantation have gained in popularity (10, 26, 27). Transplantation consists of mixing into the dysbiosed system a taxa-diverse microbiota sample, either originating from its native habitat or in some purified preparation. The aim here is to allow recolonization of the dysbiosed habitat, reset the community development path and restore a well-functioning microbiome. Although the outcomes of microbiota transplants have been astonishing (28, 29), the processes and mechanisms underlying microbiota mergers are very poorly understood (30). By producing two types of carefully controlled taxa-complex resident soil microbiota and a designed synthetic soil community of 21 native soil isolates (SynCom) in the same habitat, we were able to precisely follow community trajectories before and after merging. We show how transplanted SynCom at between 50 to 80% of the mixed cells at start is able to temporarily increase the productivity of an indigenous regrown soil community (SoilCom) as well as of a suboptimal soil-grown community (LakeCom) but is then displaced to less than 1% by members from the taxa-diverse merger communities (Fig. 3E). SynCom members were displaced regardless of their taxonomic identity. Since the soil microcosm habitats were identical for the pre-merged, merged and control community growth, we conclude that the displacement of the SynCom members – even though originating from the same native site, must be due to strain and/or functional redundancies in the more complex partner communities (SoilCom and LakeCom). This indicates that nutritional niche competition and emerging interspecific interactions select the functionalities of strains that can proliferate in the merged communities under growing conditions, and that the number of redundant strains sharing these nutritional niches then controls their eventual relative abundances. Despite their displacement to low proportions, however, the temporal presence and or activity of SynCom members was sufficient to profoundly influence the trajectories and the outcomes of the merged communities in comparison to non-transplanted regrown communities of the same starting material. Outcomes of reverse transplants between SynCom and SoilCom inoculum under non-growth conditions suggest that the proportion of viable (SynCom) cells determines their propensity to maintain dominant colonization levels in the soil microcosms, with lower viability leading to stronger invasion and compositional turnover by the transplanted community.

A key point in our study was the controlled development of soil microbiomes with inocula from two different origins that we tried to contrast as much as possible, in order to allow precise tracking of community mergers. For one inoculum, we used washed cells from a forest top-soil, but for the other, we used filter-collected freshwater lake microorganisms. Despite different source material and unique compositional successions, both communities converged to typical soil microbiota (Fig. 1F), which indicates the strong filtering role by the soil microcosm habitat (Fig. 1F). Our results thus present experimental proof for the original hypothesis of Baas-Becking, who stated that: “*Everything is everywhere, but the environment selects*” (31). He suggested that environmental selection drives between-community differences, since he assumed that microbes can easily be dispersed everywhere (31). The premise that all (microbial) species are everywhere has been challenged by high-depth sequencing characterization of microbial species distributions in different habitats, indicating that most environments tend to have specific community signatures and dispersal is effectively limiting all microbes to be everywhere (32,10, 33). However, we were surprised that freshwater lake taxa colonized the soil habitat with mature compositions being similar to what is known from global soil community signatures (Fig. 1F, Fig. 5). This demonstrates the strong role of habitat filtering in favouring the proliferation of taxa present in the lakewater inoculum under the soil microcosm conditions. Probably this worked, because the lake inoculum already contained sufficient taxa capable of growing in the soil habitat, as a result of the natural dispersal connection between soil and lake via run-off waters (23, 34, 35). This shows, *inter alia,* that freshwater lakes could serve as transplant inoculum for the regeneration of impoverished soils, which could facilitate the handling (i.e., sprinkling transplant inoculum) as opposed to digging and mixing soil transplants into soil.

Despite their global compositional similarities, LakeComs were more diverse but less productive than SoilComs in the same habitat, with 3 to 10-fold lower maximum attained community size (Fig. 1B, 1C). Quantitative taxa comparisons among both developmental trajectories suggested that the community size difference was not caused by the absence of specific families from LakeCom inoculum, but rather by underperformance of the genera within those families in the soil microcosms (Fig. 2F). This may be due to genus or strain-specific functional differences that we did not capture in our 16S rRNA gene amplicon characterization of both communities. What seems similar at family level in the freshwater lake and the soil inoculum may thus have already been selected for utilization of different nutritional niches and will not grow as effectively in the same soil habitat. Since we did not conduct metagenomic analysis of both communities, we cannot at this point describe the exact underlying strain-level functional microdiversity among lake and soil families. We cannot exclude, however, that the different taxa starting proportions in the lakewater inoculum set the community on a different growth trajectory, leading to distinct emerging interspecific interactions and resulting in less effective nutrient utilisation than accomplished by the soil inoculum (despite encoding overall similar metabolic functions). It is not unreasonable to assume that the soil inoculum members coevolved to some extent to use soil nutrients more optimally in a collective food web via metabolite leakage and cross-feeding, resulting in better community growth (36–39). A high-diverse taxa inoculum (as in LakeCom) would thus not automatically produce higher total community biomass as a result of being able to occupy more nutritional niches than a lower taxa-diverse inoculum (36, 40, 41). Depending on the emerging interspecific interactions, a higher taxa diversity may become counterproductive for community development, leading to ‘hump-shaped’ relations of richness and productivity (42–44).

Little is known about the processes underlying microbial community coalescence from merged communities (16). Some studies have defined coalescence outcomes from the overall dominance of one or the other of the starting communities in the merger, for example, asymmetrical or symmetrical (45, 46). Other studies have related the coalescence effect to the productivity of either of the starting communities to that of the merger. For example, coalesced communities of methanogenic consortia mostly resembled the more productive starting partner (47). However, we show that this is not a general rule since the more-productive SynCom was outcompeted in the same habitat by the less-productive LakeCom. Rather, our results indicate that the redundancy of functionally similar strains, guilds or taxa is a strong determinant for successful proliferation in the merger (at least under growing conditions), and therefore, for the resulting mature community. We did observe, though, some interesting differences between mergers of LakeCom or SoilCom with SynCom that point to other processes during merged growth (Fig. 3E). In particular, merged LakeCom was characterized by a temporal blooming (between day 3-7) of the Lysobacter isolate present in SynCom (Fig. 3E). Considering that Lysobacter is described as facultative predator (48, 49), its temporal blooming may be the result of predation on other community members. SynCom decline in the mergers may thus be a consequence of both cell death, predation and the inability of SynCom members to compete effectively for nutrients with (functionally redundant and more diverse) LakeCom or SoilCom members. Unfortunately, we have no specific information on the relative abundances of potential other predators (e.g., protists) or phages in LakeCom and SoilCom, which makes it difficult to estimate the roles of predatory control (apart from the Lysobacter example) on growth of the merged communities as a whole, or of specific populations present in either of the starting communities (Fig. 3D and 3E). The role of cell death and viability is also evident from the reverse transplant experiments with pre-colonised SynComs and freshly washed SoilCom inoculum. Even in that case where no new nutrients were added, SoilCom taxa managed to proliferate, establish and eventually dominate the merged community (Fig. 6E, 6G). This domination was faster and stronger with fewer viable SynCom cells in the habitat. Since we also measured a loss of the total cell fraction, this suggests dead and lysed cells to be reutilized by some SoilCom taxa. Coalescence outcomes thus probably depend on functional overlaps and differences among individual taxa, their interactions during community development, and the proportion of viable cells (45).

Our results are significant for the general behaviour of coalescing communities and set a basis for developing transplants as means to restore dysbiosed soil communities (although we acknowledge that we did not study the potential influence of plant growth). The consistent SynCom decline upon the mergers is reason for caution when considering coalescence with synthetic guilds for precise microbiome interventions (21, 50). On the other hand, even its transient presence was sufficient to consistently and profoundly change the trajectories of the merged communities (Fig. 3E, 4B, 4C). This is in agreement with previous studies that found permanent changes in defined synthetic communities even after transient inoculant survival (51, 52). The underlying process here may be metabolite leakage from the initially highly abundant SynCom members that profits growth of the taxa in the more diverse merger community, but leads to loss of competitiveness of the SynCom members, as shown for other individual inoculants (53). For soil microbiome restorations, it may thus not be sufficient to choose transplant-taxa on the basis of their origin and assume they will proliferate and integrate into the target community (at high levels). The outcomes of the coalescence may rather depend on emerging community interaction networks and functional niche redundancies. However, the strong environmental filtering by the soil habitat as observed here can help to steer transplanted merged microbiomes in the direction of natural soil microbiome types. In conclusion, our study presents a novel approach for the *de novo* generation of soil-like microbiomes and mergers, offering new perspectives for interventions in soil restoration management.

## MATERIALS AND METHODS

### Isolation and preparation of resident and transplant communities

Three communities were used in the study, which were named SoilCom, LakeCom and SynCom. SoilCom was started from inoculum recovered by washing cells from forest topsoil (-5 to -15 cm; location: Dorigny campus, University of Lausanne, Switzerland GPS 46.52126, 6.57864) as described previously (22). Briefly, large debris (e.g., rocks, branches and leaves) were manually removed and soil was immediately homogenized and sieved through a 2 mm-grid size sieve. 250 grams of sieved soil was mixed with 500 ml of tetrasodium-pyrophosphate decahydrate solution (Sigma-Aldrich, 0.2% wt/vol, pH 7.5), homogenized for 2 minutes at 2000 rpm in a Stare SM4 blender (Satrap), and left idle to sediment for 30 min. The supernatant was divided to 50 ml polypropylene centrifuge tubes and centrifuged at 800 rpm for 5 min in an Eppendorf A-4-62 Swing Bucket Rotor to remove large particles. The upper phase was collected and layered over 10 ml sterile 60% w/v Histodenz solution in water (Sigma-Aldrich, D2158, density of 1.31 g ml^-1^) in 50 ml tubes, which were centrifuged at 3220 x g for 30 min at room temperature (Eppendorf A-4-62 Swing Bucket Rotor). The interface layer containing microbial cells was recovered, diluted five times with pyrophosphate solution, and subsequently centrifuged at 3220 x g for 10 min as above. The resulting cell pellet was resuspended in 5 ml of soil buffer (containing per L, 0.6 g of MgSO_4_·7H_2_O, 0.1 g of CaCl_2_ and 1.8 mL of 5 x M9 minimal salts solution [BD Biosciences]) and an aliquot was taken for cell counting by flow cytometry (see below). The cell density was adjusted to 10^7^ cells ml^-1^ by addition of soil extract, and 10 ml of this suspension was used to inoculate soil microcosms (see below). In the case of Syn2w and Syn3m transplants with SoilCom, the SoilCom cell suspension was diluted with soil buffer to obtain a density of 10^7^ cells ml^-1^ and 1 ml was used for mixing with the SynComs in the soil microcosms.

LakeCom originated from the freshwater Lake Geneva (Lac Léman) sampled at Port de Pierrettes (Saint Sulpice, Switzerland, GPS 46.516937, 6.579464). Water was sampled (in February 2023) ca. 5 m away from the quay wall with a sterile stainless steel bucket and immediately transported to the laboratory. Microbial biomass was collected from 10 litres of water filtered through five 47 mm diameter 0.2–µm polyethersulfone membrane filters (Type 15407, Sartorius Stedim Biotech) using a sterile glass filter holder (Millipore). Microbial cells were detached from each filter by 1 min vortexing in 5 ml sterile lake water and pooled together. Sterile lake water was prepared in advance using the same sampling and filtering procedure, but keeping the filtrate, which was subsequently autoclaved for 30 min at 121°C.

The resulting lake water cell suspension was quantified with flow cytometry (see below), centrifuged for 10 min at 3220 x g (Eppendorf A-4-62 Swing Bucket Rotor) and the cell pellet was resuspended in soil extract to obtain a density of 10^7^ cells ml^−1^, which was then used for inoculation of the soil microcosms (see below).

The SynCom was assembled from 21 individual cultured soil isolates obtained from the same location and soil type as SoilCom, as described previously (22). The SynCom members belong to genera: Microbacterium, Mucilaginibacter, Curtobacterium, Variovorax, Flavobacterium, Cellulomonas, Tardiphaga, Devosia, Mesorhizobium, Burkholderia, two species of Pseudomonas, Luteibacter, Chitinophaga, Lysobacter, Rhodococcus, Caulobacter, Cohnella, Rahnella, Phenylobacterium and Bradyrhizobium. Strains were recovered from their –80°C stocks by plating on R2A (DSMZ GmbH) and incubated for one week at 23°C. Grown colonies for each of the strains were washed from the plates with 5 ml soil buffer. The turbidity of the cell suspensions was then measured using an Ultrospec 500 pro (Amersham Biosciences) and diluted to an OD_600_ of 1 by addition of soil buffer. Equal volumes of each individual SynCom member suspension were then mixed into a single mixed culture, which was centrifuged at 3220 x g (Eppendorf A-4-62 Swing Bucket Rotor) for 10 min. The supernatant was discarded, and the cell pellet was resuspended in soil extract, to be finally diluted to an OD_600_ of 0.01 (equivalent to 10^7^ cells ml^−1^).

### Soil microcosm preparation

Soil microcosms consisted of sterilised riverbank material to which a sterile aqueous soil extract (SE) was added as described previously (22). Briefly, ca. 10 kg material was sampled in the 0-10 cm horizon, transported to the laboratory and dried for two weeks at ambient air. The dried sediment was then sieved to obtain a particle fraction ranging between ca. 0.5 and 3 mm, divided into 2 kg portions and autoclaved twice for 1 h at 121°C (followed by a dry cycle), with one week storage at 23°C in between. Aliquots of 90 g of the autoclaved matrix were then transferred into 500-ml capped Schott glass bottles and autoclaved once more. To prepare SE, freshly collected forest topsoil was mixed with tap water in a 1:1 *w/v* ratio in 5 L stainless steel containers, autoclaved for 1 h at 121°C, mixed and left to sediment overnight. Subsequently, the supernatant was decanted, centrifuged at 5000 x g for 15 min to remove insoluble solids, transferred to 500-ml glass bottles and autoclaved again. Both soil matrix and SE were checked for sterility by plating on R2A agar, and were considered sterile if no colonies were detected after one week incubation at 23 °C. The mineral and nutritional composition of soil matrix and SE have been described previously (22).

The final soil microcosms contained 90 g sterile soil matrix supplemented with 10 ml SE, providing thus ca. 10% gravimetric water content. For soil-to-soil microcosm transfers, the proportions of soil matrix, SE and the inocula were adjusted accordingly to maintain constant water content (see below). Each condition was done in four replicates.

### Microcosm inoculation and coalescence designs

Sterile soil microcosms were inoculated with 10 ml of SoilCom, LakeCom or SynCom suspensions in SE to achieve starting densities of ca. 10^6^ cells g^−1^ (Fig. 1A). After inoculation, the microcosms were immediately mixed on a roller-board and incubated upright in the dark at room temperature (23 °C). Mixing was repeated before each sampling. SoilCom and LakeCom microcosms were sampled immediately at the start (T_0_), and after 3, 7, 14 and 21 days.

Microbiome coalescence was studied by combining individual communities (SoilCom, LakeCom or SynCom) grown separately for 21 days in soil microcosms (four replicates for each condition, Fig. 3A). In this case, 500-ml Schott glass bottles were filled with 80 g of sterile soil matrix and 8 ml of SE, to which either 10 g of SoilCom and 10 g of SynCom, or 10 g of LakeCom and 10 g of SynCom were mixed. As controls, 10 g of each individual community was inoculated into 80 g of sterile soil matrix with 8 ml of SE. The resulting coalescence microcosms were again mixed by rolling, incubated in the dark at 23°C, and sampled at start, and after 3, 7, 21 and 60 days.

To test reverse coalescence, SynComs were grown for two weeks (Syn2w) or three months (Syn3m), after which ca. 1 ml of fresh SoilCom inoculum (for preparation, see above) was added to achieve 10^5^ cells per gram microcosm material (Fig. 6A). No further SE-solution was added. As control, the same density of SoilCom inoculum was inoculated in the soil microcosms comprised of sterile matrix and deionised water (without SE). All microcosms were sampled immediately after inoculation and mixing, and then further after 1, 3, 7 and 28 days of incubation.

### Microcosm sampling

For sampling, 10 g of microcosm material was removed aseptically using Sterileware sampling spatulas (SP Bel-Art) and transferred into a 50-ml sterile polypropylene tube. Cells were extracted from samples by mixing with 10 ml of pyrophosphate solution (see above), vortexing for 2 min at maximum speed on a Vortex-Genie 2 (Scientific Industries, Inc.) with vertical adaptor (SI-V525) and two minutes of passive sedimentation in a tube rack. The supernatant containing the cells was transferred to a fresh 15 ml tube, from which two aliquots of 250 µl were removed for community size enumeration using colony forming unit (CFU) counting and/or flow cytometry (see below). The remainder (5-7 ml) was centrifuged at 3220 x g for 7 min in a Eppendorf A-4-62 swing bucket rotor, and the resulting cell pellet was resuspended in 2 ml soil buffer, transferred to 2-ml tubes (Eppendorf), harvested by centrifugation at 7000 x g for 7 minutes and stored at –80°C until DNA isolation (see below).

### Community size determinations

Sizes of the communities in the samples were measured by CFU counting on R2A plates or by flow cytometry. Cell suspensions from each sample were serially diluted using soil buffer and aliquots of 10 µl were dropped on R2A plates in four technical replicates. After drying, plates were incubated in the dark at room temperature. Colonies per dried droplet area were counted using a 10× magnifying binocular after two days incubation, taking only those dilutions into account with between 1–50 colonies per area. Samples from uninoculated microcosms, soil buffer (used for sample dilution) and pyrophosphate solution used for cell extraction were taken along as sterility controls.

For flow cytometry counting a 1-ml aliquot of cell suspension from each sample was filtered using a 40–µm strainer (Falcon) and mixed with an equal volume of 4 M sodium-azide solution to fix the cells (Sigma-Aldrich). Two technical replicates of 200 µl per sample each were stored at 4°C until further processing the next day. One of the technical replicates was stained with SYBR Green I for 20 min in the dark according to the Thermo Fisher protocol; the other was kept unstained. A 10–µl aliquot of each sample was then aspired on a CytoFLEX Flow Cytometer (Beckman Coulter) with 10 µl min^−1^ flow rate. The difference in the FITC-H channel signal between stained and non-stained, and the comparison to a stained sample from a non-inoculated microcosm was used to define the cell counting gate (example shown in Fig. S6).

### DNA isolation and amplicon sequencing

DNA was extracted from –80 °C-stored microcosm recovered cell samples using the DNeasy PowerSoil Pro kit (Qiagen) according to the manufacturer’s protocol. Recovered DNA concentrations were measured using a Qubit dsDNA BR kit (Invitrogen), and stored at -20°C until library preparation.

Libraries for the 16S rRNA V3-V4 amplicon sequencing were prepared as in the Illumina protocol (https://support.illumina.com/content/dam/illumina-support/documents/documentation/chemistry_documentation/16s/16s-metagenomic-library-prep-guide-15044223-b.pdf), starting from 10 ng purified DNA per sample. Libraries were indexed using the Nextera XT Index Kit v2 set A and B (Illumina). PCR products were purified using 0.85 v CleanNGS beads (CleanNA) and quantified by Qubit BR DNA assay (Thermo Fisher Scientific). Following equimolar pooling and final bead purification, the multiplexed libraries were paired-end sequenced on a MiSeq instrument using the 300-cycle MiSeq reagent kit v3 and 30% PhiX-DNA spike-in control at the University of Lausanne Genomics Technology Facility (GTF).

Sequencing data was cleaned and analysed with Qiime 2 (version: 2021.8) within a Singularity container (version 3.8.5) on UNIX. Unique amplified sequence variants (ASV) were taxonomically referenced to SILVA (version 132). Feature and taxonomy tables, reference sequences and phylogenetic tree were exported as *phyloseq* object (R package, version: 1.40.0) for diversity analysis (see below). SynCom ASVs within coalesced samples were identified based on ASVs detected in the SynCom alone controls. All identified ASVs were then collectively used to assess the SynCom fraction levels across conditions.

### MicrobeAtlas comparison

All sample sequences were compared with a global background of soil communities from the MicrobeAtlas database (https://microbeatlas.org). Raw 16S reads from all samples were standardized and quality-filtered using a custom C++ program internal to the MicrobeAtlas pipeline and mapped using MAPseq 1.2.6 (54)(16S reference database: MAPref v2.2; all other parameters kept at default) to obtain 97%-level OTU count tables compatible with MicrobeAtlas.

MicrobeAtlas was filtered for soil, lake, and sediment samples as follows. All samples with extracted meta-data annotations “soil”, “soil|plant” (main environment) or “aquatic;lake”, “aquatic;lake|sediment” (main + sub environment) were kept, with “soil|plant” typically corresponding to rhizosphere samples according to manual checks, resulting in 184,448 samples. Samples with sub-environment “lake|sediment” were labeled as “sediment” for downstream analyses. To further exclude potential mislabeled samples, we employed a conservative nearest-neighbor-based approach where only samples whose closest neighboring samples predominantly shared the same main environment annotation as the focal sample were retained (one neighboring sample per project, up to 10 projects, min. Bray-Curtis similarity 0.2). While this excluded many potentially relevant samples, it helped ensure consistency and reduce noise of our sample sets. For “lake|sediment” samples, both “soil” and “aquatic” neighbors were considered matches. We manually checked the “organism” annotation field of all remaining 119,511 samples and blacklisted problematic terms such as “phyllosphere metagenome“, “seawater metagenome” and “uncultured eukaryote”, as well as food- or host-related terms, which resulted in 109,503 samples. Finally, we removed samples that contained more than 10% eukaryotic reads to conservatively remove samples targeting the eukaryotic fraction, leaving 81,627 total samples.

### Data processing and analysis

Data processing and statistical analysis were done using MATLAB (v.2021b) and R 4.0 (R Core Team, 2019) on RStudio (version 2022.2.3.492) using the following packages: *phyloseq* (55), *microbiome* (56), *vegan* (57), *biomformat* (58), *tidyverse* (59), *dplyr* (60), *reshape2* (61), *robCompositions* (62), *PMCMRplus* (63), *pairwiseAdonis* (64), *rstatix* (65), *emmeans* (66), *MicrobiotaProcess* (67), *ggplot2* (68), *ggpubr* (69), *Biostrings* (70), *DescTools* (71), *car* (72), *multcomp* (73), and *RVAideMemoire* (74). Flow cytometry data was imported with the *fca_readfcs* (75) function on MATLAB and analysed using custom scripts (gating procedure shown in Fig. S6).

Data normality and homogeneity of variance were always checked (where applicable) using Shapiro-Wilk’s normality test and Levene’s test, respectively. Differences in community sizes of LakeCom and SoilCom during assembly and after coalescence with SynCom (CFU and flow cytometry data), differences in LakeCom and SoilCom Chao1 values over time and SynCom absolute abundances within different conditions (across time) were compared using two-way repeated measures ANOVA on ranked values, checking for the effect of community and incubation time. The presence of outliers was tested using the *identify_outliers* function (*rstatix* package). Sphericity of data was checked automatically (as a part of *anova_test* function, *rstatix* package) and corrected using Greenhouse-Geisser correction. Residuals were checked for normality and homogeneity of variance as indicated above. *Post hoc* pairwise testing was done using *t*-tests and *P*-values were always adjusted using Holm’s method. Differences of Shannon’s index values were checked in the same manner. Two-way repeated measures ANOVA was done on actual values (not on ranked ones). Community development (with LakeCom and SoilCom assembly and upon their coalescence with SynCom), was compared by sample ordination using NMDS on Bray-Curtis pairwise distances at ASV, Family or Phylum levels (level is always indicated on the figure). In all cases, PERMANOVA (with 999 iterations, performed using *adonis2* function) was used to evaluate the influence of time and community on measured distances. Data homogeneity was checked with *betadisper* (*vegan* package). Pairwise differences between samples were then checked using the *pairwise.adonis* function from the *pairwiseAdonis* package (*P*-values adjusted with Holm method). All absolute taxa abundances were calculated by multiplying the total community size (measured with either flow cytometry or CFU plating) with the relative abundance of that taxon obtained with amplicon sequencing (see above). Fast-growing genera were identified in SoilCom and LakeCom, based on their per-genus individual absolute abundance increase from day 1 to day 3. Any taxon showing increase was counted as ‘fast-growing’ and the extent of the increase in their population was computed by subtracting the day 1 abundance from the same genus abundance at day 3. Distributions of fast-growing taxa growth in LakeCom and SoilCom were compared using a Wilcoxon rank sum test (we tested both absolute abundances coming from flow cytometry and from CFU data).

Family-level absolute abundances (from CFU and flow cytometry data) per replicate over time were used to calculate the area-under-curve value (AUC) using the *AUC* function from *DescTools* package. Mean AUC values (of 4 replicates) were paired for each family between SoilCom and LakeCom, and subsets were defined as follows (without having overlaps): “Same”, same family AUC (±0.5 log_10_) value between SoilCom and LakeCom; “LakeCom-enriched” and “SoilCom-enriched”, AUC value higher in LakeCom or SoilCom, respectively, and non-zero in the other community; and finally, “SoilCom-only” and “LakeCom-only”, positive family AUC value for either SoilCom or LakeCom, respectively, and zero for the other.

ASV-level shifts due to mergers with SynCom were evaluated by pairing abundances of common ASVs found in the following comparisons: LakeCom+SynCom vs. LakeCom, SoilCom+SynCom vs. SoilCom, and LakeCom or SoilCom replicates by themselves in all possible paired combinations. The last two comparisons were used to evaluate baseline replicate-level variation in the data. SynCom ASVs were removed, and all samples were subsampled randomly to a density of 100,000 reads. Within each comparison group, ASVs were paired (e.g., value X = abundance in the non-merged community, value Y=same ASV in the merged community) and abundances were log_10_ transformed. A linear regression was calculated on the T_0_ data per comparison group, to obtain slope coefficients and outlier thresholds (defined as 99.7% confidence interval, ± 3 standard error of residual distribution, Fig. 4B and C). The T_0_ slope and thresholds were then imposed on the data from the remaining time points to detect outlier ASVs surpassing the threshold. The total outlier distance was then obtained by summing the distance of each outlier ASV to its expected value (per condition, time point and optionally per direction - enriched or depleted). Total outlier distances were compared across all time points and for all four comparison groups in a repeated measures ANOVA on ranked values (as described above). To further inspect the effect of SynCom transplants, the distribution of residuals from a comparison of non-merged vs. merged communities was compared to the residuals from the paired comparison of either LakeCom or SoilCom replicates alone (randomly subsampled to obtain equally sized data sets). In that case, the difference between conditions was tested as a difference in data dispersion using Levene’s test for homogeneity of variance (*P*-values adjusted using Holm’s method). In addition, the median values were compared with a Wilcoxon rank sum test (shown in Supplementary Data). Source tracking analysis was conducted using the FEAST R package (version 0.1.0) (84).

For the SynCom reverse transplant experiments with SoilCom inoculum, the CFU count differences between conditions (Syn2w, Syn2w+SoilCom, Syn3m, Syn3m+SoilCom) was tested at the last time point only with a one-way ANOVA (assumptions and residuals checked as before) and *post hoc t*-tests (Holm’s correction). The last time point comparison using flow cytometry data, Chao1 or Shannon index values was made in a Kruskal-Wallis rank sum test and *post hoc* Dunn test with Holm’s correction. Differences in beta diversity of each condition (Bray-Curtis distances) were checked as described above.

UMAP (76) projections were generated from Bray-Curtis distance matrices (computed on relative abundances, normalized including unmapped reads) using julia 1.9.3 (77) and the *Distances.jl* package (78) (version 0.10.11). UMAP projections were computed using the python *umap* package (79) (version 0.5.3; parameters: n_neighbors = 3000, min_dist = 1.5, spread = 15, epochs = 500). Scatterplots were produced using python 3.9.9 (80) and the *seaborn* package (81) (version 0.13.2). Outlier studies detected in the UMAP projection were labelled using the *dbscan* algorithm from the *Clustering.jl* package (82)(version 0.15.2; parameters: radius = 1.5, min_cluster_size = 5), leaving 63,899 samples that underwent a final round of UMAP projection with identical parameters.

For the differential abundance analysis, a distinct, intermediary zone (IZ) of the projection was delineated based on i) the initial trajectory of SoilCom and LakeCom samples, and ii) cluster distinctness (Fig. S1B). This resulted in 3733 IZ and 60,017 non-IZ samples. To account for sample imbalance and computational complexity, 1000 samples of each group were randomly selected for further analysis. To check for differentially abundant 97% OTUs between the two zones (IZ and non-IZ) we used SiamCat v.2.2.0 (83) in R 3.6.3. All OTUs with a prevalence of <0.5% or a maximum abundance of <0.0001% were excluded. The associations were tested with a Wilcoxon Rank-sum test, adjusted for multiple testing via the Benjamini Hochberg method. P=0.05 was used as our significance threshold. We then created a siamcat object to plot the results in an association plot, and to assess fold change of OTUs in IZ versus non-IZ.

## Acknowledgements

The authors acknowledge the support of Lausanne Genomics Technology Facility in the amplicon sequencing and Lisa Teglbjaerg for help in preliminary experiments.

## Funding

This work was supported by the Swiss National Centre in Competence Research NCCR Microbiomes (No. 51NF40_180575 and 51NF40_ 225148).

## Author contributions

SC, JT, VS and JM conceived the study and designed experiments. SC and VS carried out experiments. SC obtained community size and composition data. JT and LM performed comparisons to MicrobeAtlas database and differential abundance analysis. All authors analysed output data. SC and JM prepared the draft manuscript. JM and CM provided funding. All authors approved the final manuscript.

## Competing interests

The authors declare that they have no competing interests.

## Data and code availability

Community profiling data (v3-v4 16S rRNA amplicon sequences) is accessible through BioProject ID PRJNA1192999. All code for data processing and raw data, organized per manuscript figure, will be made available for download from Zenodo.

## Notes

### Competing Interest Statement

The authors have declared no competing interest.

