## Supplementary figures and data for "Habitat filtering but not microbiota origin controls microbiome transplant outcomes in soil"

Senka Causevic *et al.*

**This PDF file includes:**

Figs. S1 to S6
Supplementary Data

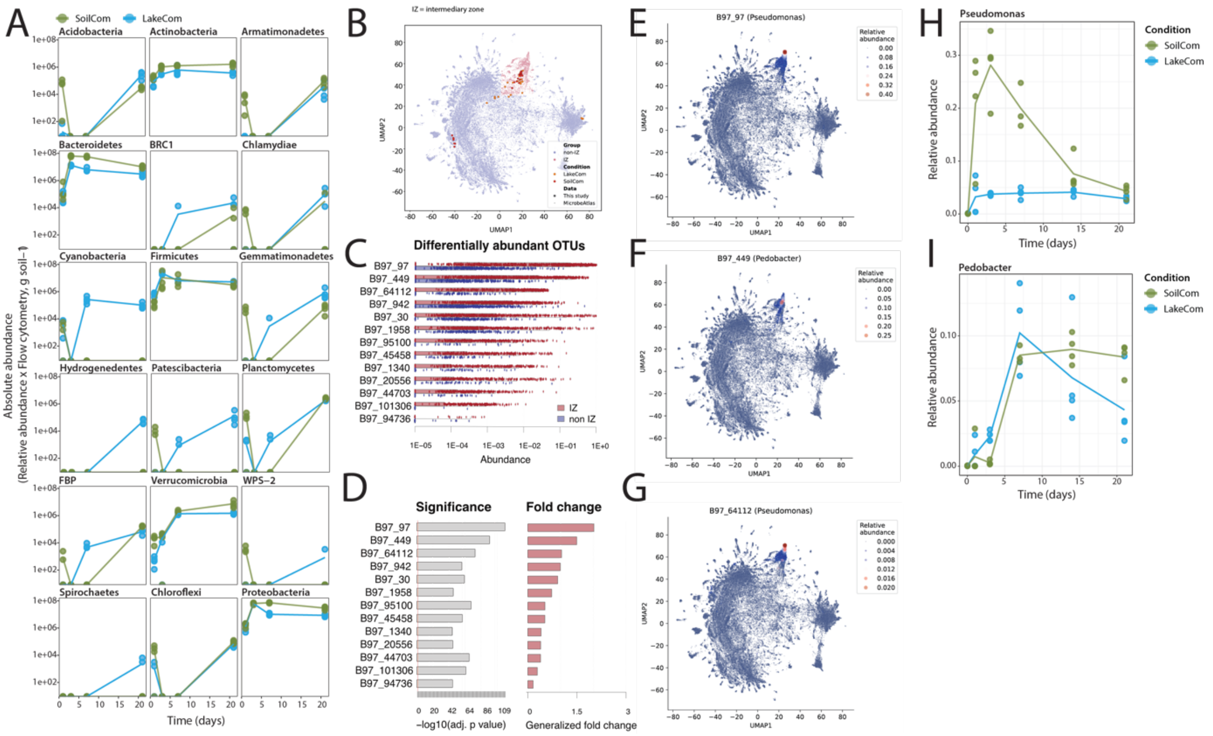


**Fig. S1. SoilCom and LakeCom compositional trajectories. (A)** Values (presented as dots, 4 biological replicates) obtained using relative abundance of each phylum and total community size measured with flow cytometry. Lines connect mean values per condition over time. Colour code as indicated on the figure. Phyla only showing a quick decline from day 0 to 1 are not included. **(B)** UMAP projection of SoilCom, LakeCom, and MicrobeAtlas samples, coloured by IZ (intermediary zone, red) and non-IZ (blue) regions. **(C)** Relative abundances of OTUs differentially abundant in 1000 randomly selected IZ (red) and non-IZ (blue) samples. The top 13 significantly enriched OTUs are plotted. **(D)** Generalized fold-chance and adjusted p-values of OTUs significantly enriched OTUs in the IZ. **(E)** UMAP projection of MicrobeAtlas samples, point color and size according to the relative abundance of the top IZ-enriched OTU B97_97 (classified: Pseudomonas). **(F)** and **(G)** As in (E), but for OTUs ranked second and third: B97_449 (Pedobacter) and OTU B97_64112 (Pseudomonas). **(H)** and **(I)** Relative abundance of Pseudomonas and Pedobacter genus, respectively, in LakeCom (in blue) and SoilCom (in green) condition. Lines connect mean of biological replicates (presented as dots).

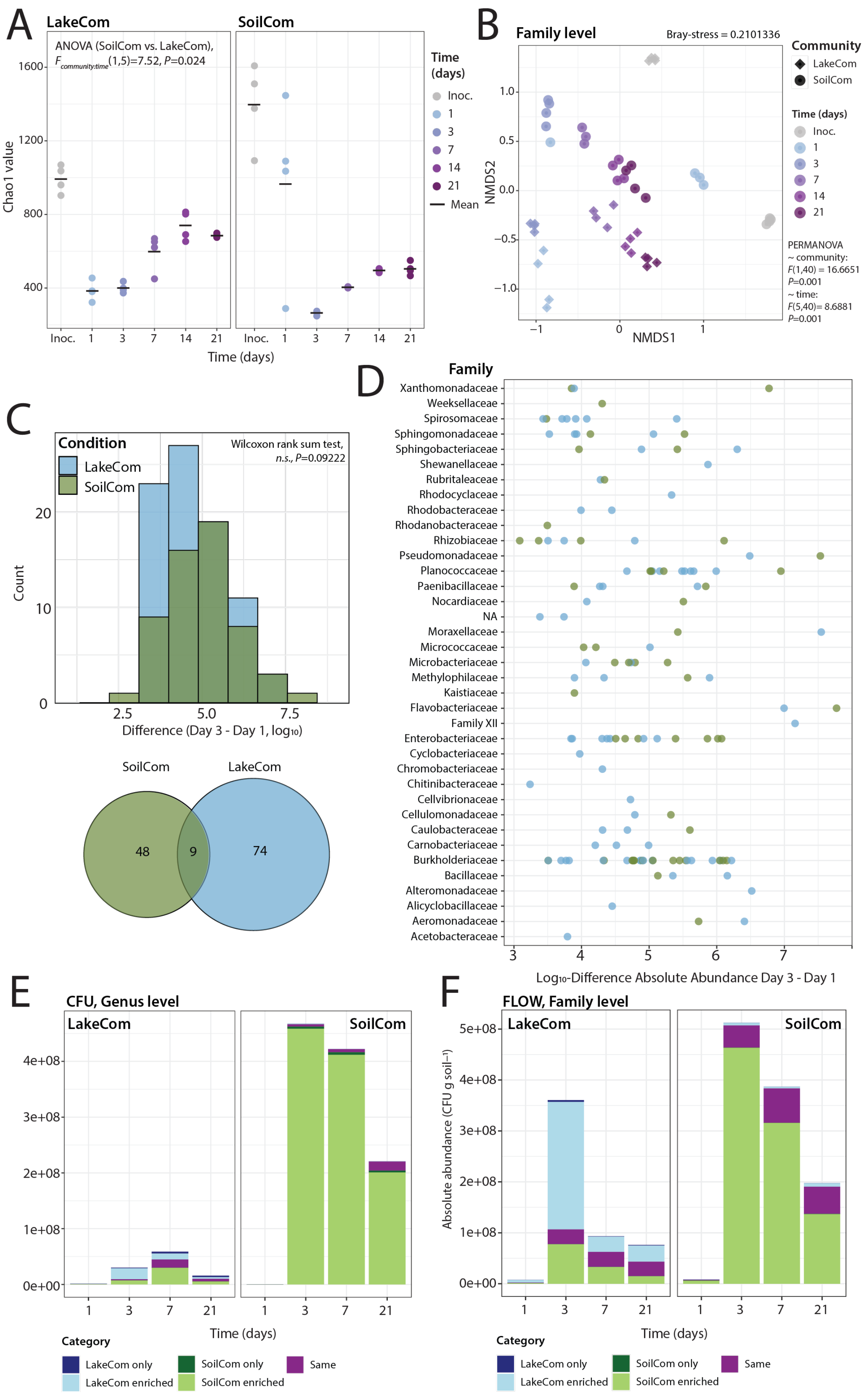
A
LakeCom
SoilCom
ANOVA (SoilCom vs. LakeCom),
$F_{community:time}$(1,5)=7.52, P=0.024
Chao1 value
1600
1200
800
400
Inoc. 1 3 7 14 21
Time (days)
Time (days)
Inoc.
1
3
7
14
21
Mean
B
Family level
Bray-stress = 0.2101336
NMDS2
NMDS1
1.0
0.5
0.0
−0.5
−1.0
−1
0
1
Community
LakeCom
SoilCom
Time (days)
Inoc.
1
3
7
14
21
PERMANOVA
~ community:
F(1,40) = 16.6651
P=0.001
~ time:
F(5,40)= 8.6881
P=0.001
C
Condition
LakeCom
SoilCom
Wilcoxon rank sum test,
n.s., P=0.09222
Count
0
10
20
2.5
5.0
7.5
Difference (Day 3 - Day 1, log10)
SoilCom
LakeCom
48
9
74
D
Family
Xanthomonadaceae
Weeksellaceae
Spirosomaceae
Sphingomonadaceae
Sphingobacteriaceae
Shewanellaceae
Rubritaleaceae
Rhodocyclaceae
Rhodobacteraceae
Rhodanobacteraceae
Rhizobiaceae
Pseudomonadaceae
Planococcaceae
Paenibacillaceae
Nocardiaceae
NA
Moraxellaceae
Micrococcaceae
Microbacteriaceae
Methylophilaceae
Kaistiaceae
Flavobacteriaceae
Family XII
Enterobacteriaceae
Cyclobacteriaceae
Chromobacteriaceae
Chitinibacteraceae
Cellvibrionaceae
Cellulomonadaceae
Caulobacteraceae
Carnobacteriaceae
Burkholderiaceae
Bacillaceae
Alteromonadaceae
Alicyclobacillaceae
Aeromonadaceae
Acetobacteraceae
3 4 5 6 7
Log10-Difference Absolute Abundance Day 3 - Day 1
E
CFU, Genus level
LakeCom
SoilCom
4e+08
3e+08
2e+08
1e+08
0e+00
1 3 7 21
Time (days)
Category
LakeCom only
LakeCom enriched
SoilCom only
SoilCom enriched
Same
F
FLOW, Family level
LakeCom
SoilCom
Absolute abundance (CFU g soil−1)
5e+08
4e+08
3e+08
2e+08
1e+08
0e+00
1 3 7 21
Time (days)
Category
LakeCom only
LakeCom enriched
SoilCom only
SoilCom enriched
Same

**Fig. S2. Changes in diversity and productivity of soil and freshwater communities (based on flow cytometry measurements). (A)** Chao1 index values (individual dots) describe richness of SoilCom and LakeCom biological replicates (4 per condition) over time. Colour gradient follows the progression of time (from grey to purple) and black lines indicate mean per condition and time point. Two-way repeated measures ANOVA on ranked values was used to evaluate effect of community type and time on measured values. **(B)** NMDS ordination based on Bray-Curtis pairwise sample distances calculated from family-level compositions. Colour code as in (A), circles and diamonds depict SoilCom and LakeCom condition, respectively. PERMANOVA with 999 iterations was used to evaluate the effect of community and time (see Materials and Methods and Supplementary Data). **(C)**,**(D)** and **(F)** as in Fig. 2C, 2D and 2F respectively, except all analysis was done using absolute abundances calculated from flow cytometry and not CFU counts. **(E)** As in 2F, except analysis was done on Genus level for CFU-inferred absolute abundances.

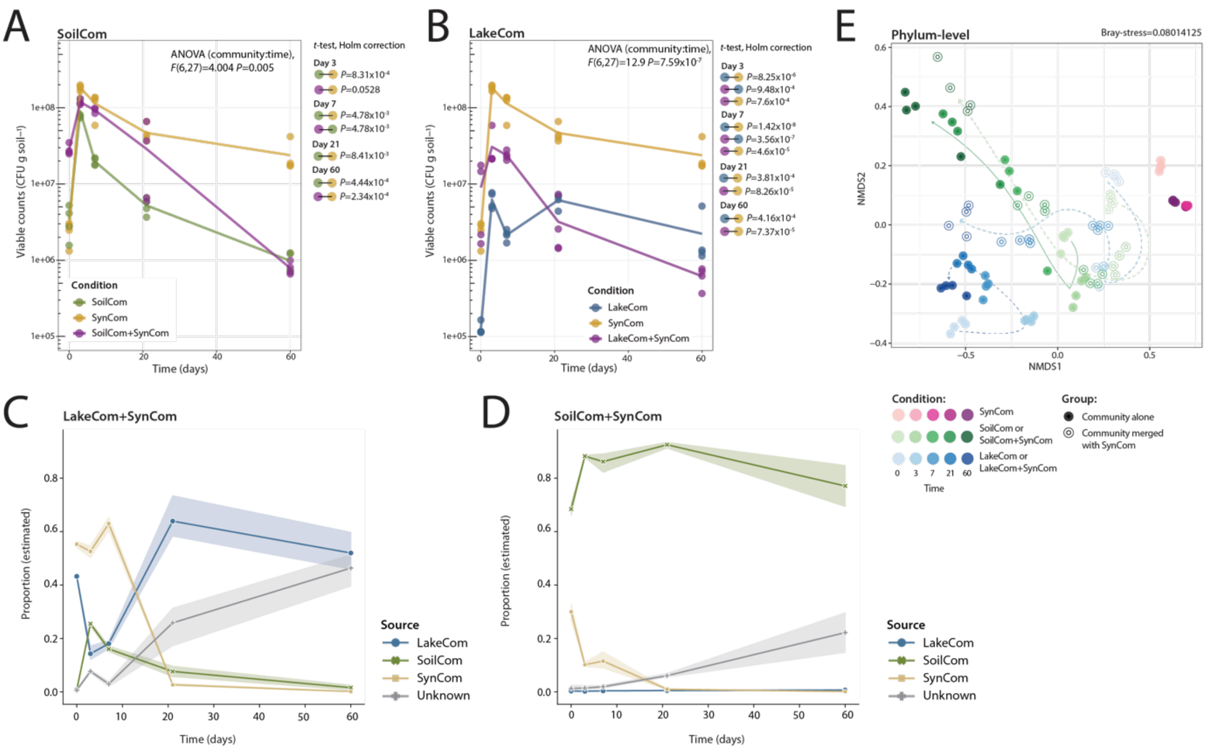


**Fig. S3. Community coalescence impacts community productivity transiently and shifts community compositional trajectories. (A)** Viable community size per gram of soil (i.e., CFU counts) is shown as dots for biological replicates of SoilCom (green), SoilCom+SynCom (magenta) and SynCom (yellow), with lines connecting mean values per time point and condition. Two-way repeated measures ANOVA on ranked values with *post hoc t*-tests (with Holm correction) was done to test for community and time effects. Only *P*-values below 0.05 are indicated on the figure with comparison groups indicated with circles coloured as described for community conditions. **(B)** As in (A), but for LakeCom (blue), LakeCom-SynCom (magenta) and SynCom (in yellow, replicated from (A) for comparison with lake conditions). **(C)** Estimated proportions of source communities (LakeCom, SoilCom, SynCom) within merged LakeCom+SynCom communities over time, as predicted by the microbial source tracking algorithm FEAST. SoilCom was included as a control. **(D)** Same as (C), but for merged SoilCom+SynCom communities. LakeCom was included as a control. **(E)** Phylum-level compositions of all merged and non-merged communities across time were compared pairwise using Bray-Curits distances and ordinated with NMDS. Shades of green, blue and magenta are used to illustrate progression of time with soil, lake and SynCom samples (respectively). Samples coming from SoilCom or LakeCom which was merged with SynCom, are depicted as simple circles, whereas circles of non-merged community samples are filled in. Arrows were added manually to increase readability and show community trajectory over time. Dashed, lighter arrows show trajectories of merged communities, and the full, darker lines are used for non-merged communities.

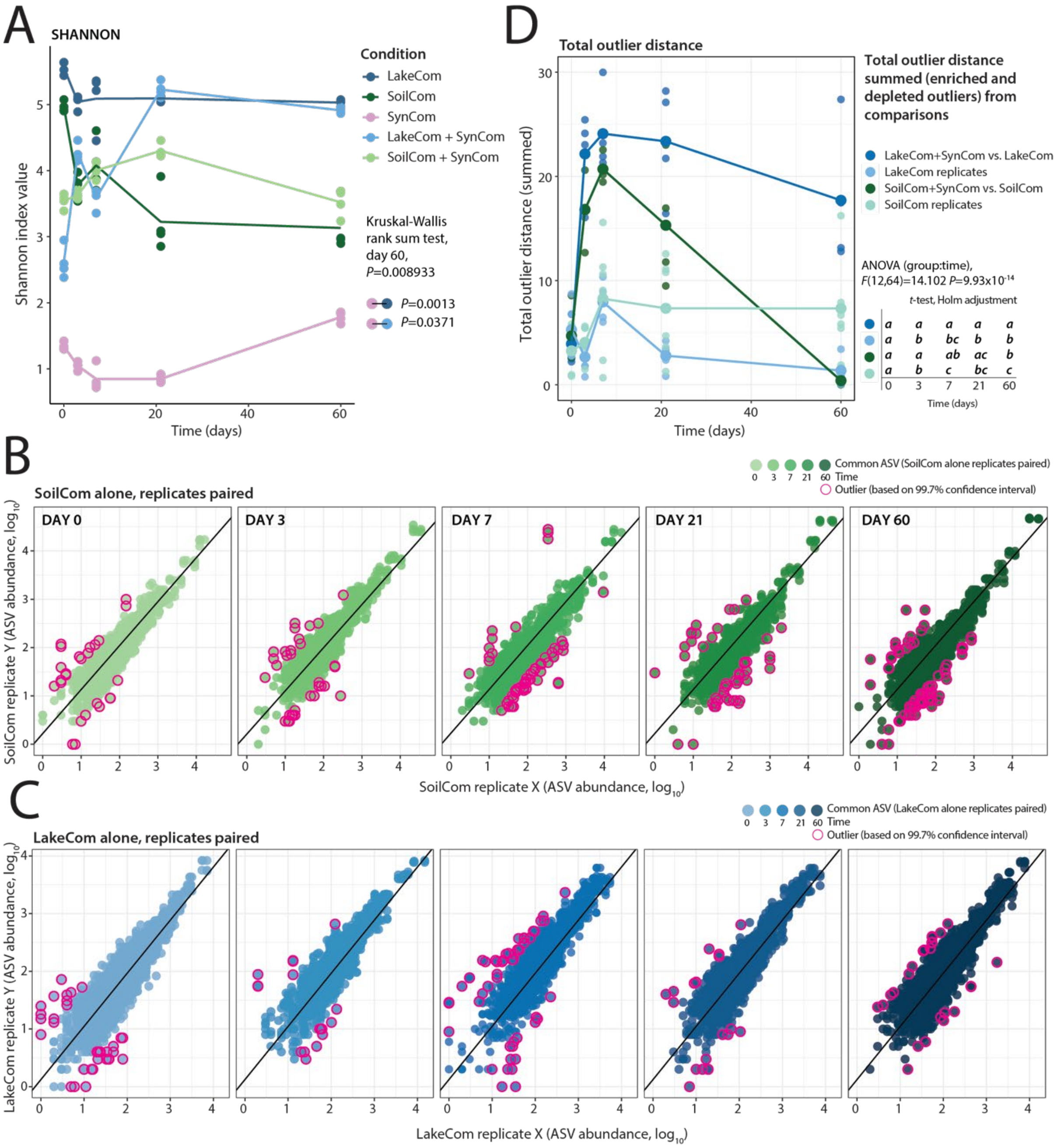


**Fig. S4. Baseline SoilCom and LakeCom taxa displacement over time. (A)** Dots show Shannon index values for all conditions (biological replicate values) with mean values over time connected with a line. Colour code included in the figure. Community effect on measured diversity levels was tested for the last time point using Kruskal-Wallis rank sum test. *Post hoc* testing was done using *t*-tests (*P*-values adjusted with Holm method). **(B)** Dots present ASV abundances paired within replicates (all possible combinations) of non-merged SoilCom (previously subsampled to 100000 reads) over time (time points presented as shades of green). Black line shows the slope obtained on $T_0$ data comparison, which is used to define outliers of all time points (based on a 99.7% confidence interval). Outliers are presented as dots with magenta circles. **(C)** As in (B), but for comparison among non-merged LakeCom replicates. **(D)** Distance of outliers (identified in between-replicate, non-merged community and merged vs. non-merged comparisons) to expected values is summed across time, per replicate (individual dots) and comparison group. Mean total distance across time is shown using lines (colours indicated on the figure). Comparison within replicates of non-merged communities contained all possible replicate combinations (in total 6), while the merged vs. non-merged comparison had 4 replicate combinations. The effect of time and comparison group was tested using two-way repeated measures ANOVA on ranked values. Letters on the right side of the figure show per-comparison similarities based on pairwise *post hoc* *t*-tests (full report in Supplementary Data).

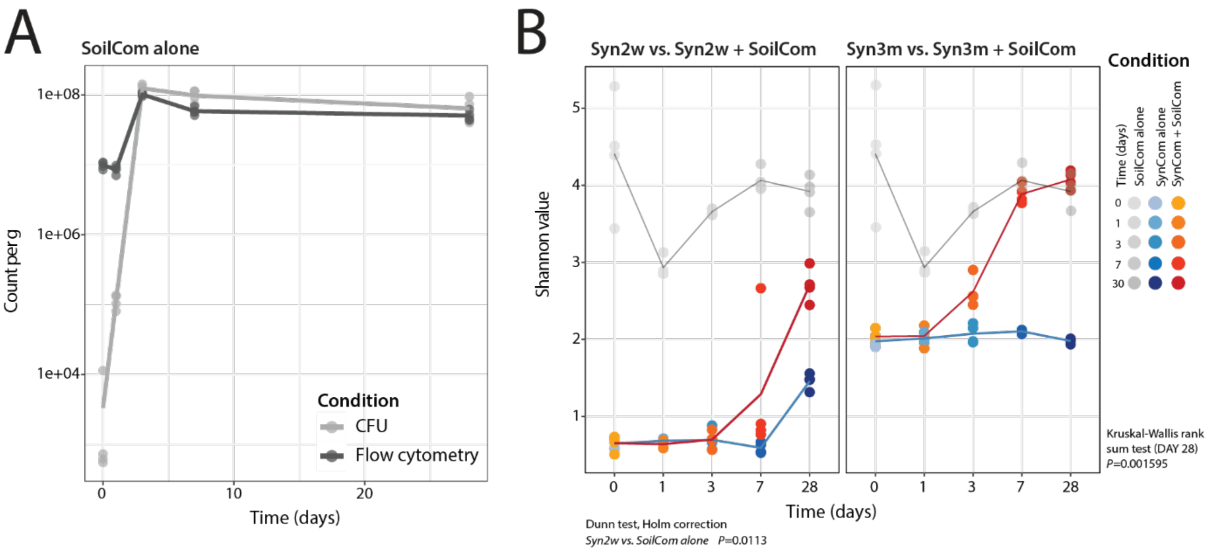


**Fig. S5. SynCom colonisation resistance upon soil cell wash inoculation. (A)** Lines connect the mean community size (of 4 biological replicates) of soil cell wash incubated in soil microcosms over time, measured with CFU (light grey) counting and flow cytometry (dark grey). **(B)** Dots show Shannon index value for each replicate of Syn2w, Syn2w+SoilCom, Syn3m, Syn3m+SoilCom and SoilCom alone over time. Lines connect the means per condition over time (note that the x-axis is discretised). Differences of Shannon's values between conditions were checked for the last time point measurement with Kruskal-Wallis rank sum test with *post hoc* Dunn test (Holm correction included). Only significant *P*-values are indicated (<0.05) on the figure.

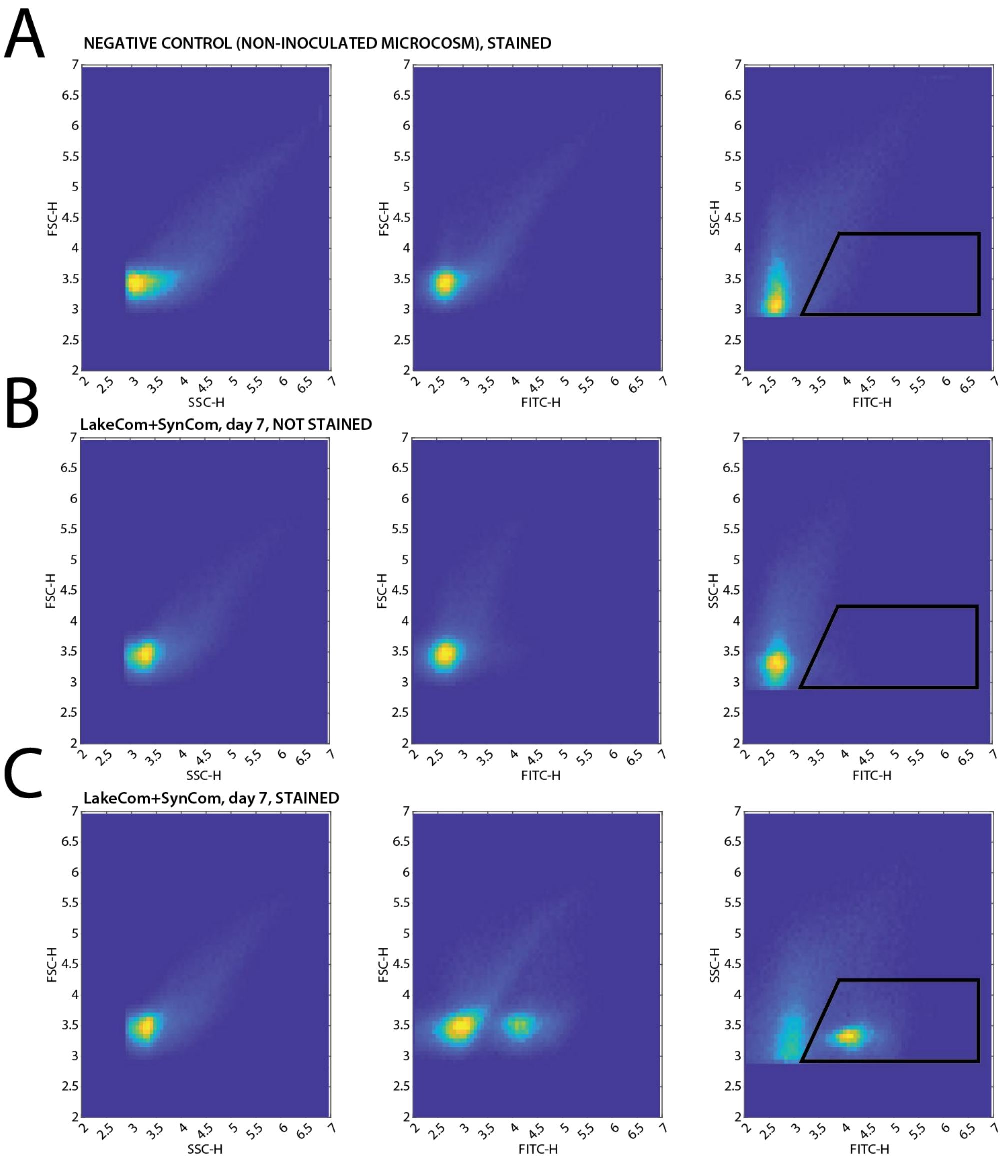


**Fig. S6. Gating strategy for community size measurement using flow cytometry data. (A)** Flow cytometry events of not inoculated microcosm example (negative control) stained with Sybr Green II. From left to right, events in FSC-H vs. SSC-H, FSC-H vs. FITC-H and SSC-H vs. FITC-H. Colour scale from blue to bright yellow (white) relates to event density. Community gate established from the last comparison as indicated on the figure. **(B)** As in (A), but for non-stained LakeCom+SynCom sample from day 3 measurement. **(C)** As in (A) and (B), but for stained LakeCom+SynCom sample.

**Supplementary Data**

### Fig. 1B

Shapiro-Wilk normality test

W = 0.75916 p-value = 7.603e-06

Levene's Test for Homogeneity of Variance (center = median)

| Df | F value | Pr(>F) |
|---|---|---|
| 7 | 3.7403 | 0.007072 |
| 24 | | |

Outliers

| Condition | Time | Sample | Phase | CFU | ranked value | is.outlier | is.extreme |
|---|---|---|---|---|---|---|---|
| LakeCom | 21d | LW-2 | 1 | 4400000 | 9 | TRUE | FALSE |
| LakeCom | 7d | LW-3 | 1 | 19000000 | 20 | TRUE | FALSE |
| SoilCom | 21d | NC2-4 | 1 | 90800000 | 24 | TRUE | FALSE |

Repeated measures two-way ANOVA with ranked values

| Effect | DFn | DFd | F | p | ges |
|---|---|---|---|---|---|
| Condition | 1 | 6 | 351.086 | 1.49E-06 | 0.902 |
| Time | 3 | 18 | 225.716 | 1.82E-14 | 0.969 |
| Condition:Time | 3 | 18 | 49.058 | 7.25E-09 | 0.873 |

Pairwise *t*-test (on ranked values), Holm's correction

| Time | .y. | group1 | group2 | n1 | n2 | p | p.adj | p.adj.signif |
|---|---|---|---|---|---|---|---|---|
| 1 | ranked value | LakeCom | SoilCom | 4 | 4 | 0.00412 | 0.00412 | ** |
| 3 | ranked value | LakeCom | SoilCom | 4 | 4 | 0.0000392 | 0.0000392 | **** |
| 7 | ranked value | LakeCom | SoilCom | 4 | 4 | 0.0000895 | 0.0000895 | **** |
| 21 | ranked value | LakeCom | SoilCom | 4 | 4 | 0.0000176 | 0.0000176 | **** |

Residuals

Shapiro-Wilk normality test

W=0.95665 p-value=0.222

Levene's Test for Homogeneity of Variance (center = median)

| Df | F value | Pr(>F) |
|---|---|---|
| 7 | 0.3624 | 0.9151 |
| 24 | | |

### Fig. 1C

Shapiro-Wilk normality test

W = 0.86432 p-value = 0.0002043

Levene's Test for Homogeneity of Variance (center = median)

| Df | F value | Pr(>F) |
|---|---|---|
| 9 | 4.7499 | 0.0005627 |
| 30 | | |

Outliers

| Community | Time | Sample | Phase | Per g cell | is.outlier | is.extreme |
|---|---|---|---|---|---|---|
| SoilCom | 3d | NC2-4 3day | 1 | 120875480 | TRUE | FALSE |

Repeated measures ANOVA with ranked values

| Effect | DFn | DFd | F | p | ges |
|---|---|---|---|---|---|
| Community | 1 | 6 | 130.39 | 2.71E-05 | 0.85 |
| Time | 4 | 24 | 179.376 | 1.67E-17 | 0.957 |
| Condition:Time | 4 | 24 | 17.362 | 8.17E-07 | 0.681 |

Pairwise *t*-test (on ranked values), Holm’s correction

| Time | .y. | group1 | group2 | n1 | n2 | p | p.adj | p.adj.signif |
|---|---|---|---|---|---|---|---|---|
| 1 | ranked value | LakeCom | SoilCom | 4 | 4 | 0.429 | 0.429 | ns |
| 3 | ranked value | LakeCom | SoilCom | 4 | 4 | 0.00208 | 0.00208 | ** |
| 7 | ranked value | LakeCom | SoilCom | 4 | 4 | 0.0000525 | 0.0000525 | **** |
| 10 | ranked value | LakeCom | SoilCom | 4 | 4 | 0.000109 | 0.000109 | *** |

Residuals

Shapiro-Wilk normality test

W=0.96666 p-value=0.2807

Levene's Test for Homogeneity of Variance (center = median)

| Df | F value | Pr(>F) |
|---|---|---|
| 9 | 0.5018 | 0.8614 |
| 30 | | |

### Fig. 2A

Shapiro-Wilk normality test

W=0.96397 p-value=0.1545

Levene's Test for Homogeneity of Variance (center = median)

| Df | F value | Pr(>F) |
|---|---|---|
| 11 | 1.8686 | 0.07901 |
| 35 | | |

Outliers

| Condition | Time | Sample | Phase | Shannon | is.outlier | is.extreme |
|---|---|---|---|---|---|---|
| LakeCom | 0 | LW1 t0 | 1 | 5.01 | TRUE | FALSE |
| LakeCom | 7 | LW1 d7 | 1 | 5.25 | TRUE | FALSE |
| SoilCom | 3 | NC2 2 d3 | 1 | 4.11 | TRUE | FALSE |

Repeated measures ANOVA

| Effect | DFn | DFd | F | p | ges |
|---|---|---|---|---|---|
| Community | 1 | 5 | 0.002 | 9.66E-01 | 7.48E-05 |
| Time | 5 | 25 | 38.776 | 5.40E-11 | 8.63E-01 |
| Community:Time | 5 | 25 | 38.077 | 6.59E-11 | 8.61E-01 |

Pairwise *t*-test (on ranked values), Holm’s correction

| Time | .y. | group1 | group2 | n1 | n2 | p | p.signif | p.adj | p.adj.signif |
|---|---|---|---|---|---|---|---|---|---|
| 0 | Shannon | LA | NC | 4 | 4 | 0.00000192 | **** | 0.00000192 | **** |

| | | | | | | | | | |
|---|---|---|---|---|---|---|---|---|---|
| 1 | Shannon | LA | NC | 4 | 4 | 0.0186 | * | 0.0186 | * |
| 3 | Shannon | LA | NC | 4 | 4 | 0.0018 | ** | 0.0018 | ** |
| 7 | Shannon | LA | NC | 4 | 3 | 0.0001 | *** | 0.0001 | *** |
| 14 | Shannon | LA | NC | 4 | 4 | 0.00036 | *** | 0.00036 | *** |
| 21 | Shannon | LA | NC | 4 | 4 | 0.00026 | *** | 0.00026 | *** |

Residuals

Shapiro-Wilk normality test

W=0.75214 p-value=0.0000004886

Levene's Test for Homogeneity of Variance (center = median)

| Df | F value | Pr(>F) |
|---|---|---|
| 11 | 0.8283 | 0.6139 |
| 30 | | |

### Fig. S2A

Shapiro-Wilk normality test

W=0.87263 p-value=0.0001076

Levene's Test for Homogeneity of Variance (center = median)

| Df | F value | Pr(>F) |
|---|---|---|
| 11 | 0.974 | 0.4871 |
| 35 | | |

Outliers

| Condition | Time | Sample | Phase | Chao1 ranked | is.outlier | is.extreme |
|---|---|---|---|---|---|---|
| LakeCom | 7 | LW1 d7 | 1 | 17 | TRUE | FALSE |
| SoilCom | 1 | NC2 1 d1 | 1 | 5 | TRUE | FALSE |

Repeated measures ANOVA on ranked values

| Effect | DFn | DFd | F | p | ges |
|---|---|---|---|---|---|
| Condition | 1 | 5 | 2.44E+00 | 1.79E-01 | 0.079 |
| Time | 1.37 | 6.83 | 2.39E+01 | 0.001 | 0.797 |
| Condition:Time | 1.37 | 6.83 | 7.52E+00 | 0.024 | 0.554 |

Pairwise t-test (on ranked values), Holm’s correction

| Time | .y. | group1 | group2 | n1 | n2 | p | p.signif | p.adj | p.adj.signif |
|---|---|---|---|---|---|---|---|---|---|
| 0 | Chao1 ranked | LA | NC | 4 | 4 | 0.0228 | * | 0.0228 | * |
| 1 | Chao1 ranked | LA | NC | 4 | 4 | 0.0522 | ns | 0.0522 | ns |
| 3 | Chao1 ranked | LA | NC | 4 | 4 | 0.00185 | ** | 0.00185 | ** |
| 7 | Chao1 ranked | LA | NC | 4 | 3 | 0.0128 | * | 0.0128 | * |
| 14 | Chao1 ranked | LA | NC | 4 | 4 | 0.000257 | *** | 0.000257 | *** |
| 21 | Chao1 ranked | LA | NC | 4 | 4 | 0.00182 | ** | 0.00182 | ** |

Residuals

Shapiro-Wilk normality test

W=0.75586 p-value=5.745e-07

Levene's Test for Homogeneity of Variance (center = median)

| Df | F value | Pr(>F) |
|---|---|---|

| | | |
|---|---|---|
| 11 | 0.6903 | 0.737 |
| 30 | | |

### Fig. 2B

Bray-stress= 0.1050865

<u>Permutation test for adonis under reduced model</u>

Number of permutation: 999

| | Df | SumOfSqs | R2 | F | Pr(>F) |
|---|---|---|---|---|---|
| group | 1 | 3.7826 | 0.21484 | 18.8865 | 0.001 |
| time | 5 | 5.8124 | 0.33014 | 5.8043 | 0.001 |
| Residual | 40 | 8.0112 | 0.45502 | | |
| Total | 46 | 17.6063 | 1 | | |

<u>ANOVA on dispersion</u>

| | Df | Sum Sq | Mean Sq | F | Pr(>F) |
|---|---|---|---|---|---|
| Groups | 11 | 0.39524 | 0.035931 | 3.7013 | 0.001494 |
| Residuals | 35 | 0.33977 | 0.009708 | | |

<u>Pairwise Adonis</u>

| | pairs | Df | SumsOfSqs | F.Model | R2 | p.value | p.adjusted | sig |
|---|---|---|---|---|---|---|---|---|
| 1 | lake 1 | vs | lake 14 | 1 | 0.91846894 | 6.199349 | 0.5081705 | 0.031 |
| 2 | lake 1 | vs | lake 21 | 1 | 1.05261242 | 7.095662 | 0.541833 | 0.046 |
| 3 | lake 1 | vs | lake 3 | 1 | 0.34994672 | 3.48112 | 0.3671634 | 0.04 |
| 4 | lake 1 | vs | lake 7 | 1 | 0.67975248 | 4.835712 | 0.4462754 | 0.031 |
| 5 | lake 1 | vs | lake 0 | 1 | 1.62725461 | 17.083929 | 0.7400789 | 0.018 |
| 6 | lake 1 | vs | soil 1 | 1 | 1.16676133 | 5.594133 | 0.4824969 | 0.028 |
| 7 | lake 1 | vs | soil 14 | 1 | 1.39763665 | 9.754961 | 0.6191676 | 0.028 |
| 8 | lake 1 | vs | soil 21 | 1 | 1.42381787 | 10.160704 | 0.628729 | 0.027 |
| 9 | lake 1 | vs | soil 3 | 1 | 1.50898304 | 14.337452 | 0.7049778 | 0.016 |
| 10 | lake 1 | vs | soil 7 | 1 | 1.28327097 | 10.500425 | 0.6774282 | 0.027 |
| 11 | lake 1 | vs | soil 0 | 1 | 1.64154815 | 14.251112 | 0.70372 | 0.031 |
| 12 | lake 14 | vs | lake 21 | 1 | 0.17189112 | 1.464494 | 0.1961947 | 0.183 |
| 13 | lake 14 | vs | lake 3 | 1 | 1.15154982 | 16.556318 | 0.7339991 | 0.024 |
| 14 | lake 14 | vs | lake 7 | 1 | 0.35072547 | 3.200175 | 0.3478385 | 0.025 |
| 15 | lake 14 | vs | lake 0 | 1 | 1.7871038 | 27.803142 | 0.8225017 | 0.038 |
| 16 | lake 14 | vs | soil 1 | 1 | 1.25916229 | 7.090069 | 0.5416373 | 0.035 |
| 17 | lake 14 | vs | soil 14 | 1 | 1.27645875 | 11.366417 | 0.6545056 | 0.023 |
| 18 | lake 14 | vs | soil 21 | 1 | 1.26715914 | 11.608668 | 0.6592587 | 0.032 |
| 19 | lake 14 | vs | soil 3 | 1 | 1.62046502 | 21.817368 | 0.7843074 | 0.025 |
| 20 | lake 14 | vs | soil 7 | 1 | 1.31104312 | 15.416226 | 0.7550968 | 0.029 |
| 21 | lake 14 | vs | soil 0 | 1 | 1.71962965 | 20.419805 | 0.7728976 | 0.024 |
| 22 | lake 21 | vs | lake 3 | 1 | 1.35378409 | 19.410849 | 0.7638804 | 0.028 |
| 23 | lake 21 | vs | lake 7 | 1 | 0.58558807 | 5.33391 | 0.4706152 | 0.028 |
| 24 | lake 21 | vs | lake 0 | 1 | 1.78882169 | 27.747766 | 0.8222105 | 0.037 |
| 25 | lake 21 | vs | soil 1 | 1 | 1.28689141 | 7.238454 | 0.5467749 | 0.026 |
| 26 | lake 21 | vs | soil 14 | 1 | 1.35107618 | 12.010519 | 0.6668613 | 0.03 |
| 27 | lake 21 | vs | soil 21 | 1 | 1.3324075 | 12.185189 | 0.6700612 | 0.034 |
| 28 | lake 21 | vs | soil 3 | 1 | 1.6515716 | 22.179383 | 0.7870784 | 0.024 |

| | | | | | | | | |
|---|---|---|---|---|---|---|---|---|
| 29 | lake 21 | vs | soil 7 | 1 | 1.35739903 | 15.918594 | 0.7609782 | 0.02 |
| 30 | lake 21 | vs | soil 0 | 1 | 1.71412834 | 20.308615 | 0.7719378 | 0.035 |
| 31 | lake 3 | vs | lake 7 | 1 | 0.86930273 | 14.028468 | 0.7004264 | 0.03 |
| 32 | lake 3 | vs | lake 0 | 1 | 1.92061157 | 115.363237 | 0.9505616 | 0.028 |
| 33 | lake 3 | vs | soil 1 | 1 | 1.42009443 | 10.926616 | 0.6455287 | 0.023 |
| 34 | lake 3 | vs | soil 14 | 1 | 1.60850994 | 24.871727 | 0.8056474 | 0.029 |
| 35 | lake 3 | vs | soil 21 | 1 | 1.6410149 | 26.671187 | 0.8163519 | 0.035 |
| 36 | lake 3 | vs | soil 3 | 1 | 1.61257403 | 60.51974 | 0.9098012 | 0.038 |
| 37 | lake 3 | vs | soil 7 | 1 | 1.37738112 | 49.38855 | 0.9080689 | 0.037 |
| 38 | lake 3 | vs | soil 0 | 1 | 1.87468727 | 51.241749 | 0.8951814 | 0.03 |
| 39 | lake 7 | vs | lake 0 | 1 | 1.80281369 | 31.800929 | 0.8412737 | 0.043 |
| 40 | lake 7 | vs | soil 1 | 1 | 1.25129139 | 7.360159 | 0.5509035 | 0.031 |
| 41 | lake 7 | vs | soil 14 | 1 | 1.21980364 | 11.648858 | 0.6600347 | 0.028 |
| 42 | lake 7 | vs | soil 21 | 1 | 1.24111163 | 12.219293 | 0.6706788 | 0.033 |
| 43 | lake 7 | vs | soil 3 | 1 | 1.58239356 | 23.72844 | 0.7981731 | 0.028 |
| 44 | lake 7 | vs | soil 7 | 1 | 1.2045885 | 15.862512 | 0.7603357 | 0.026 |
| 45 | lake 7 | vs | soil 0 | 1 | 1.74489289 | 22.771145 | 0.7914577 | 0.024 |
| 46 | lake 0 | vs | soil 1 | 1 | 1.61813152 | 12.977226 | 0.6838316 | 0.024 |
| 47 | lake 0 | vs | soil 14 | 1 | 1.82103108 | 30.659269 | 0.8363306 | 0.029 |
| 48 | lake 0 | vs | soil 21 | 1 | 1.83050977 | 32.541714 | 0.8443245 | 0.024 |
| 49 | lake 0 | vs | soil 3 | 1 | 1.93554034 | 90.57714 | 0.9378735 | 0.032 |
| 50 | lake 0 | vs | soil 7 | 1 | 1.65715568 | 76.873447 | 0.9389301 | 0.033 |
| 51 | lake 0 | vs | soil 0 | 1 | 1.89762853 | 60.610257 | 0.9099238 | 0.031 |
| 52 | soil 1 | vs | soil 14 | 1 | 0.89968898 | 5.209128 | 0.464722 | 0.033 |
| 53 | soil 1 | vs | soil 21 | 1 | 1.03454705 | 6.101027 | 0.5041743 | 0.034 |
| 54 | soil 1 | vs | soil 3 | 1 | 0.7043301 | 5.229379 | 0.4656873 | 0.028 |
| 55 | soil 1 | vs | soil 7 | 1 | 0.62797162 | 3.986141 | 0.4435877 | 0.058 |
| 56 | soil 1 | vs | soil 0 | 1 | 1.03380381 | 7.148076 | 0.5436595 | 0.026 |
| 57 | soil 14 | vs | soil 21 | 1 | 0.08188759 | 0.785304 | 0.115736 | 0.713 |
| 58 | soil 14 | vs | soil 3 | 1 | 1.09200672 | 15.736597 | 0.7239678 | 0.031 |
| 59 | soil 14 | vs | soil 7 | 1 | 0.49712566 | 6.277985 | 0.5566584 | 0.026 |
| 60 | soil 14 | vs | soil 0 | 1 | 1.7097253 | 21.551373 | 0.782225 | 0.02 |
| 61 | soil 21 | vs | soil 3 | 1 | 1.30177148 | 19.64991 | 0.7660811 | 0.028 |
| 62 | soil 21 | vs | soil 7 | 1 | 0.6946835 | 9.21184 | 0.6481807 | 0.029 |
| 63 | soil 21 | vs | soil 0 | 1 | 1.71646096 | 22.529298 | 0.7896899 | 0.024 |
| 64 | soil 3 | vs | soil 7 | 1 | 0.15351121 | 4.575134 | 0.4778141 | 0.025 |
| 65 | soil 3 | vs | soil 0 | 1 | 1.83557513 | 44.438741 | 0.8810438 | 0.028 |
| 66 | soil 7 | vs | soil 0 | 1 | 1.56299104 | 34.36576 | 0.8729861 | 0.024 |

### Fig. S2B

Bray-stress=0.2101336

PERMANOVA (999 iterations)

| | Df | SumOfSqs | R2 | F | Pr(>F) |
|---|---|---|---|---|---|
| group | 1 | 1.9795 | 0.16648 | 16.6651 | 0.001 |
| time | 5 | 5.16 | 0.43395 | 8.6881 | 0.001 |
| Residual | 40 | 4.7513 | 0.39958 | | |
| Total | 46 | 11.8909 | 1 | | |

ANOVA on dispersion

| | Df | Sum Sq | Mean Sq | F | Pr(>F) |
|---|---|---|---|---|---|
| Groups | 11 | 0.28742 | 0.026129 | 2.2081 | 0.03718 |
| Residuals | 35 | 0.41417 | 0.011833 | | |

Pairwise Adonis

| | pairs | Df | SumsOfSqs | F.Model | R2 | p.value | p.adjusted | sig | |
|---|---|---|---|---|---|---|---|---|---|
| 1 | lake 1 | vs | lake 14 | 1 | 0.77269388 | 9.601507 | 0.6154218 | 0.029 | 1 |
| 2 | lake 1 | vs | lake 21 | 1 | 0.91248784 | 11.79355 | 0.6627992 | 0.036 | 1 |
| 3 | lake 1 | vs | lake 3 | 1 | 0.1592125 | 2.480326 | 0.29248 | 0.18 | 1 |
| 4 | lake 1 | vs | lake 7 | 1 | 0.53826247 | 7.18045 | 0.5447803 | 0.032 | 1 |
| 5 | lake 1 | vs | lake 0 | 1 | 1.50774645 | 23.539856 | 0.7968846 | 0.032 | 1 |
| 6 | lake 1 | vs | soil 1 | 1 | 0.95198146 | 6.481576 | 0.5192915 | 0.026 | 1 |
| 7 | lake 1 | vs | soil 14 | 1 | 0.99566221 | 12.691137 | 0.6789922 | 0.028 | 1 |
| 8 | lake 1 | vs | soil 21 | 1 | 0.98742666 | 12.823675 | 0.6812525 | 0.034 | 1 |
| 9 | lake 1 | vs | soil 3 | 1 | 1.20569841 | 16.764381 | 0.7364304 | 0.029 | 1 |
| 10 | lake 1 | vs | soil 7 | 1 | 0.9540407 | 11.839198 | 0.7030737 | 0.022 | 1 |
| 11 | lake 1 | vs | soil 0 | 1 | 1.59757245 | 24.883907 | 0.8057241 | 0.033 | 1 |
| 12 | lake 14 | vs | lake 21 | 1 | 0.12216345 | 3.675659 | 0.3798872 | 0.029 | 1 |
| 13 | lake 14 | vs | lake 3 | 1 | 0.91850575 | 45.801204 | 0.8841726 | 0.028 | 1 |
| 14 | lake 14 | vs | lake 7 | 1 | 0.19511649 | 6.329558 | 0.5133645 | 0.034 | 1 |
| 15 | lake 14 | vs | lake 0 | 1 | 1.41929945 | 71.26851 | 0.9223487 | 0.04 | 1 |
| 16 | lake 14 | vs | soil 1 | 1 | 0.86864132 | 8.454834 | 0.584914 | 0.028 | 1 |
| 17 | lake 14 | vs | soil 14 | 1 | 0.49961868 | 14.558769 | 0.7081537 | 0.023 | 1 |
| 18 | lake 14 | vs | soil 21 | 1 | 0.46492338 | 14.146755 | 0.7021853 | 0.037 | 1 |
| 19 | lake 14 | vs | soil 3 | 1 | 1.27840607 | 46.011867 | 0.8846417 | 0.031 | 1 |
| 20 | lake 14 | vs | soil 7 | 1 | 0.75040617 | 27.168895 | 0.8445704 | 0.026 | 1 |
| 21 | lake 14 | vs | soil 0 | 1 | 1.25718604 | 62.655501 | 0.9126071 | 0.029 | 1 |
| 22 | lake 21 | vs | lake 3 | 1 | 1.19428827 | 70.460973 | 0.9215286 | 0.035 | 1 |
| 23 | lake 21 | vs | lake 7 | 1 | 0.43050434 | 15.529505 | 0.7213127 | 0.039 | 1 |
| 24 | lake 21 | vs | lake 0 | 1 | 1.50937841 | 89.789026 | 0.9373623 | 0.027 | 1 |
| 25 | lake 21 | vs | soil 1 | 1 | 0.85831793 | 8.614668 | 0.5894536 | 0.025 | 1 |
| 26 | lake 21 | vs | soil 14 | 1 | 0.74242573 | 23.785916 | 0.7985625 | 0.031 | 1 |
| 27 | lake 21 | vs | soil 21 | 1 | 0.65633002 | 22.054269 | 0.7861288 | 0.033 | 1 |
| 28 | lake 21 | vs | soil 3 | 1 | 1.38611331 | 56.164045 | 0.9034812 | 0.029 | 1 |
| 29 | lake 21 | vs | soil 7 | 1 | 0.91616092 | 38.341759 | 0.8846378 | 0.034 | 1 |
| 30 | lake 21 | vs | soil 0 | 1 | 0.97333538 | 57.388322 | 0.9053453 | 0.03 | 1 |
| 31 | lake 3 | vs | lake 7 | 1 | 0.62895541 | 43.256639 | 0.878189 | 0.033 | 1 |
| 32 | lake 3 | vs | lake 0 | 1 | 1.31795732 | 363.207379 | 0.983749 | 0.032 | 1 |
| 33 | lake 3 | vs | soil 1 | 1 | 1.17087668 | 13.543528 | 0.692993 | 0.02 | 1 |
| 34 | lake 3 | vs | soil 14 | 1 | 1.18978459 | 65.984711 | 0.916649 | 0.03 | 1 |
| 35 | lake 3 | vs | soil 21 | 1 | 1.21564282 | 73.327976 | 0.9243646 | 0.031 | 1 |
| 36 | lake 3 | vs | soil 3 | 1 | 1.12361619 | 97.721786 | 0.9421529 | 0.033 | 1 |
| 37 | lake 3 | vs | soil 7 | 1 | 0.86254206 | 106.794377 | 0.955275 | 0.029 | 1 |
| 38 | lake 3 | vs | soil 0 | 1 | 1.80712933 | 478.215784 | 0.9876088 | 0.03 | 1 |
| 39 | lake 7 | vs | lake 0 | 1 | 1.17683816 | 81.720759 | 0.9316011 | 0.031 | 1 |
| 40 | lake 7 | vs | soil 1 | 1 | 0.92060788 | 9.468847 | 0.6121236 | 0.026 | 1 |
| 41 | lake 7 | vs | soil 14 | 1 | 0.36438493 | 12.650815 | 0.6782982 | 0.036 | 1 |
| 42 | lake 7 | vs | soil 21 | 1 | 0.3952681 | 14.452099 | 0.7066316 | 0.034 | 1 |

| | | | | | | | | | |
|---|---|---|---|---|---|---|---|---|---|
| 43 | lake 7 | vs | soil 3 | 1 | 1.12929374 | 50.708795 | 0.8941963 | 0.03 | 1 |
| 44 | lake 7 | vs | soil 7 | 1 | 0.5001244 | 23.811892 | 0.8264605 | 0.03 | 1 |
| 45 | lake 7 | vs | soil 0 | 1 | 1.57850344 | 108.481045 | 0.9475896 | 0.032 | 1 |
| 46 | lake 0 | vs | soil 1 | 1 | 1.30333264 | 15.099988 | 0.7156396 | 0.026 | 1 |
| 47 | lake 0 | vs | soil 14 | 1 | 1.16338761 | 65.023323 | 0.9155207 | 0.036 | 1 |
| 48 | lake 0 | vs | soil 21 | 1 | 1.16760314 | 71.027301 | 0.9221055 | 0.021 | 1 |
| 49 | lake 0 | vs | soil 3 | 1 | 1.36206275 | 119.913101 | 0.9523481 | 0.027 | 1 |
| 50 | lake 0 | vs | soil 7 | 1 | 0.92137115 | 116.490299 | 0.9588445 | 0.028 | 1 |
| 51 | lake 0 | vs | soil 0 | 1 | 1.53899151 | 422.853957 | 0.9860092 | 0.029 | 1 |
| 52 | soil 1 | vs | soil 14 | 1 | 0.7114963 | 7.064379 | 0.5407359 | 0.023 | 1 |
| 53 | soil 1 | vs | soil 21 | 1 | 0.75524493 | 7.608525 | 0.5590999 | 0.024 | 1 |
| 54 | soil 1 | vs | soil 3 | 1 | 0.47493775 | 5.042715 | 0.4566554 | 0.115 | 1 |
| 55 | soil 1 | vs | soil 7 | 1 | 0.43521776 | 4.056142 | 0.4478885 | 0.123 | 1 |
| 56 | soil 1 | vs | soil 0 | 1 | 0.72108843 | 8.33978 | 0.5815835 | 0.03 | 1 |
| 57 | soil 14 | vs | soil 21 | 1 | 0.03899249 | 1.264293 | 0.1740421 | 0.355 | 1 |
| 58 | soil 14 | vs | soil 3 | 1 | 1.03182221 | 40.053187 | 0.8697159 | 0.028 | 1 |
| 59 | soil 14 | vs | soil 7 | 1 | 0.35244553 | 13.990105 | 0.736705 | 0.026 | 1 |
| 60 | soil 14 | vs | soil 0 | 1 | 1.4987662 | 83.070556 | 0.9326377 | 0.028 | 1 |
| 61 | soil 21 | vs | soil 3 | 1 | 1.22680445 | 50.468659 | 0.8937464 | 0.034 | 1 |
| 62 | soil 21 | vs | soil 7 | 1 | 0.48747452 | 20.788874 | 0.8061179 | 0.022 | 1 |
| 63 | soil 21 | vs | soil 0 | 1 | 1.30650008 | 78.756872 | 0.9292093 | 0.038 | 1 |
| 64 | soil 3 | vs | soil 7 | 1 | 0.1354281 | 7.804411 | 0.6095096 | 0.022 | 1 |
| 65 | soil 3 | vs | soil 0 | 1 | 1.76724886 | 153.553848 | 0.9623951 | 0.037 | 1 |
| 66 | soil 7 | vs | soil 0 | 1 | 1.43550607 | 177.448462 | 0.972595 | 0.034 | 1 |

### Fig. 3B

Shapiro-Wilk normality test

W=0.94458  p-value=0.02443

Levene's Test for Homogeneity of Variance (center = median)

| Df | F value | Pr(>F) |
|---|---|---|
| 11 | 0.8268 | 0.6148 |
| 36 | | |

Outliers

| Condition | Time | Sample | Phase | Per g cell | is.outlier | is.extreme |
|---|---|---|---|---|---|---|
| SoilCom+SynCom | 7 | SYN+SOIL2 stained | 2 | 6220000 | TRUE | FALSE |

Repeated measures ANOVA on ranked values

| Effect | DFn | DFd | F | p |
|---|---|---|---|---|
| Condition | 2 | 9 | 10.867 | 4.00E-03 |
| Time | 3 | 27 | 470.791 | 2.06E-23 |
| Condition:Time | 6 | 27 | 13.645 | 4.45E-07 |

Pairwise *t*-test (on ranked values), Holm's correction

| Time | .y. | group1 | group2 | n1 | n2 | p | p.adj | p.adj.signif |
|---|---|---|---|---|---|---|---|---|
| 3 | ranked value | soilcom | syncom | 4 | 4 | 0.427 | 0.427 | ns |
| 3 | ranked value | soilcom | soilcom+syncom | 4 | 4 | 0.0479 | 0.0958 | ns |
| 3 | ranked value | syncom | soilcom+syncom | 4 | 4 | 0.0123 | 0.0369 | * |

| | | | | | | | | |
|---|---|---|---|---|---|---|---|---|
| 7 | ranked value | soilcom | syncom | 4 | 4 | 0.382 | 0.382 | ns |
| 7 | ranked value | soilcom | soilcom+syncom | 4 | 4 | 0.0195 | 0.0586 | ns |
| 7 | ranked value | syncom | soilcom+syncom | 4 | 4 | 0.0876 | 0.175 | ns |
| 21 | ranked value | soilcom | syncom | 4 | 4 | 0.000515 | 1.03E-03 | ** |
| 21 | ranked value | soilcom | soilcom+syncom | 4 | 4 | 0.0043 | 4.30E-03 | ** |
| 21 | ranked value | syncom | soilcom+syncom | 4 | 4 | 0.00000812 | 2.44E-05 | **** |
| 60 | ranked value | soilcom | syncom | 4 | 4 | 0.00000493 | 1.48E-05 | **** |
| 60 | ranked value | soilcom | soilcom+syncom | 4 | 4 | 0.000199 | 3.98E-04 | *** |
| 60 | ranked value | syncom | soilcom+syncom | 4 | 4 | 0.00568 | 5.68E-03 | ** |

Residuals

Shapiro-Wilk normality test

W=0.9592 p-value=0.09373

Levene's Test for Homogeneity of Variance (center = median)

| Df | F value | Pr(>F) |
|---|---|---|
| 11 | 0.8268 | 0.6148 |
| 36 | | |

### Fig. 3C

Shapiro-Wilk normality test

W=0.76985 p-value=2.883e-07

Levene's Test for Homogeneity of Variance (center = median)

| Df | F value | Pr(>F) |
|---|---|---|
| 11 | 0.9754 | 0.4854 |
| 36 | | |

Outliers

| Condition | Time | Sample | Phase | Per g cell | is.outlier | is.extreme |
|---|---|---|---|---|---|---|
| LakeCom | 21 | LA4 stained | 2 | 23500000 | TRUE | FALSE |
| SynCom | 21 | SA3 stained | 2 | 19600000 | TRUE | FALSE |
| SynCom+LakeCom | 21 | SL4 stained | 2 | 18900000 | TRUE | FALSE |

Repeated measures two-way ANOVA on ranked values

| Effect | DFn | DFd | F | p | ges |
|---|---|---|---|---|---|
| Condition | 2 | 9 | 1.87E+01 | 6.20E-04 | 0.717 |
| Time | 3 | 27 | 5.11E+01 | 2.91E-11 | 0.689 |
| Condition:Time | 6 | 27 | 5.54E+01 | 6.52E-14 | 0.828 |

Pairwise *t*-test (on ranked values), Holm's correction

| Time | .y. | group1 | group2 | n1 | n2 | p | p.adj | p.adj.signif |
|---|---|---|---|---|---|---|---|---|
| 3 | ranked value | lakecom | syncom | 4 | 4 | 1.32E-07 | 3.95E-07 | **** |
| 3 | ranked value | lakecom | lake+syn | 4 | 4 | 0.00000261 | 5.22E-06 | **** |
| 3 | ranked value | syncom | lake+syn | 4 | 4 | 0.00185 | 1.85E-03 | ** |
| 7 | ranked value | lakecom | syncom | 4 | 4 | 8.01E-08 | 2.40E-07 | **** |
| 7 | ranked value | lakecom | lake+syn | 4 | 4 | 0.00000021 | 4.20E-07 | **** |
| 7 | ranked value | syncom | lake+syn | 4 | 4 | 0.135 | 0.135 | ns |
| 21 | ranked value | lakecom | syncom | 4 | 4 | 0.527 | 1 | ns |

| | | | | | | | | |
|---|---|---|---|---|---|---|---|---|
| 21 | ranked value | lakecom | lake+syn | 4 | 4 | 0.844 | 1 | ns |
| 21 | ranked value | syncom | lake+syn | 4 | 4 | 0.412 | 1 | ns |
| 60 | ranked value | lakecom | syncom | 4 | 4 | 0.00186 | 0.00372 | ** |
| 60 | ranked value | lakecom | lake+syn | 4 | 4 | 0.572 | 0.572 | ns |
| 60 | ranked value | syncom | lake+syn | 4 | 4 | 0.000811 | 0.00243 | ** |

Residuals

Shapiro-Wilk normality test

W=0.97191 p-value=0.3

Levene's Test for Homogeneity of Variance (center = median)

| Df | F value | Pr(>F) |
|---|---|---|
| 11 | 0.9754 | 0.4854 |
| 36 | | |

### Fig. S3A

Shapiro-Wilk normality test

W=0.87329 p-value=0.0001123

Levene's Test for Homogeneity of Variance (center = median)

| Df | F value | Pr(>F) |
|---|---|---|
| 11 | 1.114 | 0.3791 |
| 36 | | |

Outliers

| Condition | Time | Sample | Phase | CFU | is.outlier | is.extreme |
|---|---|---|---|---|---|---|
| SoilCom | 3 | NA1 | 2 | 128000000 | TRUE | FALSE |
| SoilCom+SynCom | 7 | SN4 | 2 | NA | TRUE | FALSE |

Repeated measures two-way ANOVA on ranked values

| Effect | DFn | DFd | F | p | ges |
|---|---|---|---|---|---|
| Condition | 2.00E+00 | 9.00E+00 | 44.769 | 2.10E-05 | 0.706 |
| Time | 3.00E+00 | 27 | 103.882 | 6.09E-15 | 0.897 |
| Condition:Time | 6.00E+00 | 27 | 4.004 | 0.005 | 0.403 |

Pairwise *t*-test (on ranked values), Holm's correction

| Time | .y. | group1 | group2 | n1 | n2 | p | p.adj | p.adj.signif |
|---|---|---|---|---|---|---|---|---|
| 3 | ranked value | soilcom | syncom | 4 | 4 | 0.000277 | 0.000831 | *** |
| 3 | ranked value | soilcom | syn+soil | 4 | 4 | 0.0528 | 0.0528 | ns |
| 3 | ranked value | syncom | syn+soil | 4 | 4 | 0.00652 | 0.013 | * |
| 7 | ranked value | soilcom | syncom | 4 | 4 | 0.00159 | 4.78E-03 | ** |
| 7 | ranked value | soilcom | syn+soil | 4 | 4 | 0.00174 | 4.78E-03 | ** |
| 7 | ranked value | syncom | syn+soil | 4 | 4 | 0.954 | 9.54E-01 | ns |
| 21 | ranked value | soilcom | syncom | 4 | 4 | 0.0028 | 8.41E-03 | ** |
| 21 | ranked value | soilcom | syn+soil | 4 | 4 | 0.0482 | 9.64E-02 | ns |
| 21 | ranked value | syncom | syn+soil | 4 | 4 | 0.108 | 1.08E-01 | ns |
| 60 | ranked value | soilcom | syncom | 4 | 4 | 0.000222 | 4.44E-04 | *** |
| 60 | ranked value | soilcom | syn+soil | 4 | 4 | 0.399 | 3.99E-01 | ns |
| 60 | ranked value | syncom | syn+soil | 4 | 4 | 0.000078 | 2.34E-04 | *** |

Residuals

Shapiro-Wilk normality test

W=0.96674 p-value=0.1885

Levene's Test for Homogeneity of Variance (center = median)

| Df | F value | Pr(>F) |
|---|---|---|
| 11 | 1.114 | 0.3791 |
| 36 | | |

### Fig. S3B

Shapiro-Wilk normality test

W=0.66839 p-value=3.826e-09

Levene's Test for Homogeneity of Variance (center = median)

| Df | F value | Pr(>F) |
|---|---|---|
| 11 | 0.7403 | 0.6936 |
| 36 | | |

Outliers

| Condition | Time | Sample | Phase | CFU | is.outlier | is.extreme |
|---|---|---|---|---|---|---|
| LakeCom | 60 | LA4 | 2 | 5130000 | TRUE | FALSE |
| SynCom | 60 | SA1 | 2 | 41800000 | TRUE | FALSE |
| LakeCom+SynCom | 3 | SL2 | 2 | 58700000 | TRUE | FALSE |

Repeated measures two-way ANOVA on ranked values

| Effect | DFn | DFd | F | p | ges |
|---|---|---|---|---|---|
| Condition | 2.00E+00 | 9.00E+00 | 1.45E+02 | 1.42E-07 | 0.906 |
| Time | 3 | 2.70E+01 | 6.21E+01 | 3.03E-12 | 0.829 |
| Condition:Time | 6 | 2.70E+01 | 1.29E+01 | 7.59E-07 | 0.668 |

Pairwise *t*-test (on ranked values), Holm's correction

| Time | .y. | group1 | group2 | n1 | n2 | p | p.signif | p.adj | p.adj.signif |
|---|---|---|---|---|---|---|---|---|---|
| 3 | ranked value | lakecom | syncom | 4 | 4 | 0.00000275 | **** | 8.25E-06 | **** |
| 3 | ranked value | lakecom | syn+lake | 4 | 4 | 0.000948 | *** | 9.48E-04 | *** |
| 3 | ranked value | syncom | syn+lake | 4 | 4 | 0.00038 | *** | 7.60E-04 | *** |
| 7 | ranked value | lakecom | syncom | 4 | 4 | 4.75E-09 | **** | 1.42E-08 | **** |
| 7 | ranked value | lakecom | syn+lake | 4 | 4 | 1.78E-07 | **** | 3.56E-07 | **** |
| 7 | ranked value | syncom | syn+lake | 4 | 4 | 0.000046 | **** | 4.60E-05 | **** |
| 21 | ranked value | lakecom | syncom | 4 | 4 | 0.000191 | *** | 3.81E-04 | *** |
| 21 | ranked value | lakecom | syn+lake | 4 | 4 | 0.117 | ns | 1.17E-01 | ns |
| 21 | ranked value | syncom | syn+lake | 4 | 4 | 0.0000275 | **** | 8.26E-05 | **** |
| 60 | ranked value | lakecom | syncom | 4 | 4 | 0.000208 | *** | 4.16E-04 | *** |
| 60 | ranked value | lakecom | syn+lake | 4 | 4 | 0.0874 | ns | 8.74E-02 | ns |
| 60 | ranked value | syncom | syn+lake | 4 | 4 | 0.0000246 | **** | 7.37E-05 | **** |

Residuals

Shapiro-Wilk normality test

W=0.91223 p-value=0.001599

Levene's Test for Homogeneity of Variance (center = median)

| Df | F value | Pr(>F) |
|---|---|---|
| 11 | 0.7403 | 0.6936 |
| 36 | | |

### Fig. 3D

Shapiro-Wilk normality test

W=0.49623 p-value=5.149e-15

Levene's Test for Homogeneity of Variance (center = median)

| Df | F value | Pr(>F) |
|---|---|---|
| 19 | 1.2213 | 0.272 |
| 60 | | |

Outliers

| Condition | Time | Sample | Phase | Ranked value | is.outlier | is.extreme |
|---|---|---|---|---|---|---|
| LakeCom | 3 | LA3 | 2 | 30 | TRUE | FALSE |
| LakeCom | 60 | LA2 | 2 | 22 | TRUE | FALSE |
| SoilCom | 60 | NA3 | 2 | 8 | TRUE | FALSE |
| LakeCom SynCom | 3 | SA2 | 2 | 64 | TRUE | FALSE |

Repeated measures two-way ANOVA on ranked values

| Effect | DFn | DFd | F | p | ges |
|---|---|---|---|---|---|
| Condition | 4 | 15 | 346.736 | 1.40E-14 | 0.951 |
| Time | 3 | 45 | 71.274 | 3.99E-17 | 0.79 |
| Condition:Time | 12 | 45 | 8.477 | 4.45E-08 | 0.641 |

Pairwise *t*-test (on ranked values), Holm's correction

| Time | .y. | group1 | group2 | n1 | n2 | p | p.adj | p.adj.signif |
|---|---|---|---|---|---|---|---|---|
| 3 | ranked value | lakecom | soilcom | 4 | 4 | 3.30E-06 | 1.98E-05 | **** |
| 3 | ranked value | lakecom | syncom | 4 | 4 | 2.40E-10 | 2.40E-09 | **** |
| 3 | ranked value | soilcom | syncom | 4 | 4 | 1.56E-06 | 1.09E-05 | **** |
| 3 | ranked value | lakecom | syn+lake | 4 | 4 | 8.08E-08 | 6.46E-07 | **** |
| 3 | ranked value | soilcom | syn+lake | 4 | 4 | 2.54E-02 | 5.08E-02 | ns |
| 3 | ranked value | syncom | syn+lake | 4 | 4 | 1.22E-04 | 6.08E-04 | *** |
| 3 | ranked value | lakecom | syn+soil | 4 | 4 | 4.06E-08 | 3.65E-07 | **** |
| 3 | ranked value | soilcom | syn+soil | 4 | 4 | 8.94E-03 | 2.68E-02 | * |
| 3 | ranked value | syncom | syn+soil | 4 | 4 | 3.35E-04 | 1.34E-03 | ** |
| 3 | ranked value | syn+lake | syn+soil | 4 | 4 | 6.11E-01 | 6.11E-01 | ns |
| 7 | ranked value | lakecom | soilcom | 4 | 4 | 1.94E-05 | 1.17E-04 | *** |
| 7 | ranked value | lakecom | syncom | 4 | 4 | 5.00E-10 | 5.00E-09 | **** |
| 7 | ranked value | soilcom | syncom | 4 | 4 | 1.01E-06 | 7.05E-06 | **** |
| 7 | ranked value | lakecom | syn+lake | 4 | 4 | 1.58E-08 | 1.26E-07 | **** |
| 7 | ranked value | soilcom | syn+lake | 4 | 4 | 2.42E-04 | 9.69E-04 | *** |
| 7 | ranked value | syncom | syn+lake | 4 | 4 | 7.08E-03 | 2.12E-02 | * |
| 7 | ranked value | lakecom | syn+soil | 4 | 4 | 1.29E-08 | 1.16E-07 | **** |
| 7 | ranked value | soilcom | syn+soil | 4 | 4 | 1.77E-04 | 8.84E-04 | *** |
| 7 | ranked value | syncom | syn+soil | 4 | 4 | 9.83E-03 | 2.12E-02 | * |
| 7 | ranked value | syn+lake | syn+soil | 4 | 4 | 8.74E-01 | 8.74E-01 | ns |

| | | | | | | | | |
|---|---|---|---|---|---|---|---|---|
| 21 | ranked value | lakecom | soilcom | 4 | 4 | 4.37E-08 | 1.75E-07 | **** |
| 21 | ranked value | lakecom | syncom | 4 | 4 | 2.15E-14 | 2.15E-13 | **** |
| 21 | ranked value | soilcom | syncom | 4 | 4 | 1.42E-11 | 1.27E-10 | **** |
| 21 | ranked value | lakecom | syn+lake | 4 | 4 | 9.12E-10 | 5.47E-09 | **** |
| 21 | ranked value | soilcom | syn+lake | 4 | 4 | 4.57E-03 | 9.15E-03 | ** |
| 21 | ranked value | syncom | syn+lake | 4 | 4 | 2.59E-10 | 1.81E-09 | **** |
| 21 | ranked value | lakecom | syn+soil | 4 | 4 | 5.39E-11 | 4.31E-10 | **** |
| 21 | ranked value | soilcom | syn+soil | 4 | 4 | 1.39E-05 | 4.16E-05 | **** |
| 21 | ranked value | syncom | syn+soil | 4 | 4 | 6.03E-09 | 3.02E-08 | **** |
| 21 | ranked value | syn+lake | syn+soil | 4 | 4 | 9.25E-03 | 9.25E-03 | ** |
| 60 | ranked value | lakecom | soilcom | 4 | 4 | 2.35E-01 | 3.11E-01 | ns |
| 60 | ranked value | lakecom | syncom | 4 | 4 | 1.91E-11 | 1.91E-10 | **** |
| 60 | ranked value | soilcom | syncom | 4 | 4 | 5.39E-11 | 4.85E-10 | **** |
| 60 | ranked value | lakecom | syn+lake | 4 | 4 | 1.13E-02 | 4.51E-02 | * |
| 60 | ranked value | soilcom | syn+lake | 4 | 4 | 1.20E-01 | 3.11E-01 | ns |
| 60 | ranked value | syncom | syn+lake | 4 | 4 | 2.42E-10 | 1.94E-09 | **** |
| 60 | ranked value | lakecom | syn+soil | 4 | 4 | 3.34E-04 | 2.00E-03 | ** |
| 60 | ranked value | soilcom | syn+soil | 4 | 4 | 4.10E-03 | 2.05E-02 | * |
| 60 | ranked value | syncom | syn+soil | 4 | 4 | 1.39E-09 | 9.70E-09 | **** |
| 60 | ranked value | syn+lake | syn+soil | 4 | 4 | 1.04E-01 | 3.11E-01 | ns |

<u>Residuals</u>

<u>Shapiro-Wilk normality test</u>

W=0.91858 p-value=2.834e-07

<u>Levene's Test for Homogeneity of Variance (center = median)</u>

| Df | F value | Pr(>F) |
|---|---|---|
| 19 | 1.2213 | 0.272 |
| 60 | | |

### Fig. 3G

Bray-stress=0.08014125

<u>PERMANOVA (999 iterations)</u>

| | Df | SumOfSqs | R2 | F | Pr(>F) |
|---|---|---|---|---|---|
| Community | 4 | 4.679 | 0.58973 | 34.139 | 0.001 |
| Residual | 95 | 3.2551 | 0.41027 | | |
| Total | 95 | 7.934 | 1 | | |

<u>ANOVA on dispersion</u>

| | Df | Sum Sq | Mean Sq | F | Pr(>F) |
|---|---|---|---|---|---|
| Groups | 4 | 0.436 | 0.109001 | 13.838 | 6.25E-09 |
| Residuals | 95 | 0.74832 | 0.007877 | | |

<u>Pairwise Adonis</u>

| Pairs | Df | SumsOfSqs | F.Model | R2 | p.value | p.adjusted | sig |
|---|---|---|---|---|---|---|---|
| lake vs. soil | 1 | 0.7411684 | 21.772756 | 0.36425886 | 0.001 | 0.01 | * |
| lake vs. syncom | 2 | 3.2753281 | 176.102028 | 0.90493419 | 0.001 | 0.01 | * |
| lake vs. syn+lake | 1 | 0.8615435 | 29.582951 | 0.43772802 | 0.001 | 0.01 | * |
| lake vs. syn+soil | 1 | 1.1197684 | 32.29128 | 0.4593924 | 0.001 | 0.01 | * |

| | | | | | | | |
|---|---|---|---|---|---|---|---|
| soil vs. syncom | 1 | 2.369485 | 81.880679 | 0.68301814 | 0.001 | 0.01 | * |
| soil vs. syn+lake | 1 | 0.3801043 | 7.756209 | 0.16951162 | 0.002 | 0.01 | * |
| soil vs. syn+soil | 1 | 0.2956392 | 5.418548 | 0.124798 | 0.015 | 0.03 | . |
| syncom vs. syn+lake | 1 | 1.219696 | 50.778039 | 0.57196622 | 0.001 | 0.01 | * |
| syncom vs. syn+soil | 1 | 1.2819416 | 43.346493 | 0.53286247 | 0.001 | 0.01 | * |

### Fig. 4A

Bray-stress=0.1755407

PERMANOVA (999 iterations)

| | Df | SumOfSqs | R2 | F | Pr(>F) |
|---|---|---|---|---|---|
| Community | 3 | 11.7986 | 0.46495 | 35.033 | 0.001 |
| Residual | 4 | 5.4945 | 0.21652 | 12.236 | 0.001 |
| Total | 72 | 8.0827 | 0.31852 | | |

ANOVA on dispersion

| | Df | Sum Sq | Mean Sq | F | Pr(>F) |
|---|---|---|---|---|---|
| Groups | 19 | 0.054292 | 0.0028575 | 0.886 | 0.6005 |
| Residuals | 60 | 0.193516 | 0.0032253 | | |

Pairwise Adonis

| Pairs | Df | SumOfSqs | F.Model | R2 | p.value | p.adjusted | sig |
|---|---|---|---|---|---|---|---|
| lake vs. soil | 1 | 6.030949 | 47.35656 | 0.554809 | 0.001 | 0.006 | * |
| lake vs. syn+lake | 1 | 2.118814 | 11.3259 | 0.229614 | 0.001 | 0.006 | * |
| lake vs. syn+soil | 1 | 5.792344 | 37.15734 | 0.494394 | 0.001 | 0.006 | * |
| soilvs. syn+lake | 1 | 4.958208 | 24.61776 | 0.393143 | 0.001 | 0.006 | * |
| soil vs. syn+soil | 1 | 0.805358 | 4.731339 | 0.110723 | 0.002 | 0.006 | * |
| syn+lake vs. syn+soil | 1 | 3.891512 | 16.92383 | 0.308133 | 0.001 | 0.006 | * |

### Fig. S4A

Shapiro-Wilk normality test

W=0.86317 p-value=0.008933

Kruskal-Wallis rank sum test

Chi-squared=17.729 df=4 p-value=0.001394

Dunn test, Holm’s adjustment

| | lake | soil | syncom | syn+lake |
|---|---|---|---|---|
| soil | 0.0573 | - | - | - |
| syncom | 0.0013 | 1 | - | - |
| syn+lake | 1 | 0.4985 | 0.0371 | - |
| syn+soil | 0.2553 | 1 | 0.4985 | 1 |

### Fig. 4D

| day | test type | Lake ctrl | dataset2 | statistic | p value | p value adjusted |
|---|---|---|---|---|---|---|
| 3 | Levene's Test | Lake ctrl | Lake+Syn | 281.89974 | 6.49E-59 | 3.89E-58 |
| 3 | Levene's Test | Lake ctrl | Soil ctrl | 4.3526389 | 0.03711331 | 0.07422663 |

| | | | | | | |
|---|---|---|---|---|---|---|
| 3 | Levene's Test | Lake ctrl | Soil+Syn | 247.879104 | 4.35E-52 | 2.17E-51 |
| 3 | Levene's Test | Lake+Syn | Soil ctrl | 186.236975 | 3.31E-40 | 1.33E-39 |
| 3 | Levene's Test | Lake+Syn | Soil+Syn | 1.12130096 | 0.28979154 | 0.28979154 |
| 3 | Levene's Test | Soil ctrl | Soil+Syn | 162.145736 | 2.98E-35 | 8.93E-35 |
| 3 | WilcoxonTest | Lake ctrl | Lake+Syn | 516782 | 2.52E-21 | 1.26E-20 |
| 3 | PairwiseWilcoxon | Lake ctrl | Soil ctrl | 340327 | 1.17E-05 | 2.35E-05 |
| 3 | PairwiseWilcoxon | Lake ctrl | Soil+Syn | 430430 | 3.16E-27 | 1.89E-26 |
| 3 | PairwiseWilcoxon | Lake+Syn | Soil ctrl | 263331 | 5.66E-09 | 1.70E-08 |
| 3 | PairwiseWilcoxon | Lake+Syn | Soil+Syn | 361107 | 0.11181456 | 0.11181456 |
| 3 | PairwiseWilcoxon | Soil ctrl | Soil+Syn | 309001 | 7.51E-13 | 3.00E-12 |
| 0 | Levene's Test | Lake ctrl | Lake+Syn | 6.51508426 | 0.01075161 | 0.03082406 |
| 0 | Levene's Test | Lake ctrl | Soil ctrl | 21.6695017 | 3.40E-06 | 1.70E-05 |
| 0 | Levene's Test | Lake ctrl | Soil+Syn | 0.07970038 | 0.77772594 | 0.77772594 |
| 0 | Levene's Test | Lake+Syn | Soil ctrl | 46.4324229 | 1.21E-11 | 7.23E-11 |
| 0 | Levene's Test | Lake+Syn | Soil+Syn | 6.59733545 | 0.01027469 | 0.03082406 |
| 0 | Levene's Test | Soil ctrl | Soil+Syn | 15.7315679 | 7.52E-05 | 0.00030095 |
| 0 | PairwiseWilcoxon | Lake ctrl | Lake+Syn | 896183 | 0.35951765 | 1 |
| 0 | PairwiseWilcoxon | Lake ctrl | Soil ctrl | 842160 | 0.9784643 | 1 |
| 0 | PairwiseWilcoxon | Lake ctrl | Soil+Syn | 871225 | 0.6532616 | 1 |
| 0 | PairwiseWilcoxon | Lake+Syn | Soil ctrl | 653211 | 0.34270643 | 1 |
| 0 | PairwiseWilcoxon | Lake+Syn | Soil+Syn | 677436 | 0.64455848 | 1 |
| 0 | PairwiseWilcoxon | Soil ctrl | Soil+Syn | 661779 | 0.74181984 | 1 |
| 60 | Levene's Test | Lake ctrl | Lake+Syn | 250.838893 | 2.00E-53 | 1.20E-52 |
| 60 | Levene's Test | Lake ctrl | Soil ctrl | 32.1031074 | 1.72E-08 | 3.44E-08 |
| 60 | Levene's Test | Lake ctrl | Soil+Syn | 145.362181 | 5.45E-32 | 2.72E-31 |
| 60 | Levene's Test | Lake+Syn | Soil ctrl | 83.2048632 | 1.95E-19 | 7.81E-19 |
| 60 | Levene's Test | Lake+Syn | Soil+Syn | 3.75850519 | 0.05271674 | 0.05271674 |
| 60 | Levene's Test | Soil ctrl | Soil+Syn | 38.8292366 | 6.45E-10 | 1.93E-09 |
| 60 | PairwiseWilcoxon | Lake ctrl | Lake+Syn | 707191 | 1.07E-34 | 4.29E-34 |
| 60 | PairwiseWilcoxon | Lake ctrl | Soil ctrl | 411464 | 2.37E-16 | 4.74E-16 |
| 60 | PairwiseWilcoxon | Lake ctrl | Soil+Syn | 154044 | 9.12E-30 | 2.74E-29 |
| 60 | PairwiseWilcoxon | Lake+Syn | Soil ctrl | 326376 | 1.79E-05 | 1.79E-05 |
| 60 | PairwiseWilcoxon | Lake+Syn | Soil+Syn | 133210 | 1.98E-58 | 1.19E-57 |
| 60 | PairwiseWilcoxon | Soil ctrl | Soil+Syn | 83286 | 4.29E-48 | 2.15E-47 |
| 7 | Levene's Test | Lake ctrl | Lake+Syn | 171.370345 | 3.01E-37 | 1.51E-36 |
| 7 | Levene's Test | Lake ctrl | Soil ctrl | 0.0289012 | 0.86502889 | 0.86502889 |
| 7 | Levene's Test | Lake ctrl | Soil+Syn | 106.81414 | 2.97E-24 | 8.91E-24 |
| 7 | Levene's Test | Lake+Syn | Soil ctrl | 184.996865 | 5.54E-40 | 3.32E-39 |
| 7 | Levene's Test | Lake+Syn | Soil+Syn | 9.02625477 | 0.00270346 | 0.00540691 |
| 7 | Levene's Test | Soil ctrl | Soil+Syn | 115.300395 | 5.15E-26 | 2.06E-25 |
| 7 | PairwiseWilcoxon | Lake ctrl | Lake+Syn | 376475 | 7.62E-13 | 3.81E-12 |
| 7 | PairwiseWilcoxon | Lake ctrl | Soil ctrl | 428533 | 2.49E-32 | 1.49E-31 |
| 7 | PairwiseWilcoxon | Lake ctrl | Soil+Syn | 357023 | 7.85E-12 | 3.14E-11 |
| 7 | PairwiseWilcoxon | Lake+Syn | Soil ctrl | 341127 | 0.29756547 | 0.59513095 |
| 7 | PairwiseWilcoxon | Lake+Syn | Soil+Syn | 301942 | 0.47701135 | 0.59513095 |
| 7 | PairwiseWilcoxon | Soil ctrl | Soil+Syn | 296619 | 0.03426334 | 0.10279002 |
| 21 | Levene's Test | Lake ctrl | Lake+Syn | 299.522231 | 6.22E-63 | 3.11E-62 |
| 21 | Levene's Test | Lake ctrl | Soil ctrl | 33.8908293 | 7.07E-09 | 1.41E-08 |
| 21 | Levene's Test | Lake ctrl | Soil+Syn | 317.098373 | 3.46E-65 | 2.07E-64 |

| | | | | | | |
|---|---|---|---|---|---|---|
| 21 | Levene's Test | Lake+Syn | Soil ctrl | 72.1962211 | 4.25E-17 | 1.28E-16 |
| 21 | Levene's Test | Lake+Syn | Soil+Syn | 1.35983753 | 0.24371834 | 0.24371834 |
| 21 | Levene's Test | Soil ctrl | Soil+Syn | 81.246727 | 7.53E-19 | 3.01E-18 |
| 21 | PairwiseWilcoxon | Lake ctrl | Lake+Syn | 763190 | 1.96E-37 | 1.17E-36 |
| 21 | PairwiseWilcoxon | Lake ctrl | Soil ctrl | 344766 | 2.52E-18 | 1.26E-17 |
| 21 | PairwiseWilcoxon | Lake ctrl | Soil+Syn | 353472 | 0.72992798 | 0.72992798 |
| 21 | PairwiseWilcoxon | Lake+Syn | Soil ctrl | 264570 | 3.20E-05 | 6.41E-05 |
| 21 | PairwiseWilcoxon | Lake+Syn | Soil+Syn | 297286 | 1.44E-17 | 5.74E-17 |
| 21 | PairwiseWilcoxon | Soil ctrl | Soil+Syn | 154680 | 3.26E-06 | 9.79E-06 |

#### Fig. 6B

Only last day compared

Shapiro-Wilk normality test

W=0.94829 p-value=0.4631

Levene's Test for Homogeneity of Variance (center = median)

| Df | F value | Pr(>F) |
|---|---|---|
| 3 | 2.7876 | 0.08619 |
| 12 | | |

Outliers

| Condition | Time | Sample | CFU g | is.outlier | is.extreme |
|---|---|---|---|---|---|
| Syn2w | 28 | Ni 16 | 58700000 | TRUE | FALSE |
| Syn3m | 28 | Ni 8 | 11000000 | TRUE | FALSE |

ANOVA

| | Df | SumSq | MeanSq | F value | Pr(>F) |
|---|---|---|---|---|---|
| Condition | 3 | 1.96E+15 | 6.54E+14 | 6.324 | 0.0081 |
| Residuals | 12 | 1.24E+15 | 1.03E+14 | | |

Pairwise *t*-test, Holm's correction

| .y. | group1 | group2 | n1 | n2 | p | p.signif | p.adj | p.adj.signif |
|---|---|---|---|---|---|---|---|---|
| CFU g | SoilCom+Syn2w | SoilCom+Syn3m | 4 | 4 | 0.53 | ns | 1 | ns |
| CFU g | SoilCom+Syn2w | Syn2w | 4 | 4 | 0.223 | ns | 0.668 | ns |
| CFU g | SoilCom+Syn3m | Syn2w | 4 | 4 | 0.534 | ns | 1 | ns |
| CFU g | SoilCom+Syn2w | Syn3m | 4 | 4 | 0.0175 | * | 0.0699 | ns |
| CFU g | SoilCom+Syn3m | Syn3m | 4 | 4 | 0.00527 | ** | 0.0263 | * |
| CFU g | Syn2w | Syn3m | 4 | 4 | 0.00164 | ** | 0.00984 | ** |

Residuals

Shapiro-Wilk normality test

W=0.98116 p-value=0.9721

Levene's Test for Homogeneity of Variance (center = median)

| Df | F value | Pr(>F) |
|---|---|---|
| 3 | 2.7876 | 0.08619 |
| 12 | | |

### Fig. 6C

Shapiro-Wilk normality test

W=0.80334 p-value=0.003018

Kruskal-Wallis rank sum test

chi-squared=11.912 df=3 p-value=0.007692

Dunn test, Holm's adjustment

| | SoilCom+Syn2w | SoilCom+Syn3m | Syn2w |
|---|---|---|---|
| SoilCom+Syn3m | 0.3069 | - | - |
| Syn2w | 0.6559 | 0.1879 | - |
| Syn3m | 0.2988 | 0.0038 | 0.3626 |

### Fig. 6D

Shapiro-Wilk normality test

W=0.64327 p-value=1.56e-14

Kruskal-Wallis rank sum test

chi-squared=17.857 df=4 p-value=0.001316

Dunn test, Holm's adjustment

| | SoilCom | Syn2w | SoilCom+Syn2w |
|---|---|---|---|
| Syn2w | 0.0021 | - | - |
| SoilCom+Syn2w | 0.3836 | 0.3836 | - |
| Syn3m | 0.0371 | 1 | 1 |

### Fig. S6A

Only last day compared

Shapiro-Wilk normality test

W=0.91968 p-value=9.404e-06

Kruskal-Wallis rank sum test

chi-squared=17.429 df=4 p-value=0.001595

Dunn test, Holm's adjustment

| | SoilCom | Syn2w | SoilCom+Syn2w |
|---|---|---|---|
| Syn2w | 0.0113 | - | - |
| SoilCom+Syn2w | 0.7544 | 0.335 | - |
| Syn3m | 0.1621 | 1 | 1 |

### Fig. 6F

Bray-stress= 0.2337854

Permutation test for adonis under reduced model

Number of permutation: 999

| | Df | SumOfSqs | R2 | F | Pr(>F) |
|---|---|---|---|---|---|
| condition | 4 | 17.351 | 0.59408 | 34.759 | 0.001 |

| | | | | |
|---|---|---|---|---|
| Residual | 95 | 11.856 | | 0.40592 |
| Total | 95 | 29.207 | | 1 |

ANOVA on dispersion

| | Df | Sum Sq | Mean Sq | F | Pr(>F) |
|---|---|---|---|---|---|
| Groups | 4 | 2.7729 | 0.69322 | 17.426 | 9.40E-11 |
| Residuals | 95 | 3.7793 | 0.03978 | | |

Pairwise Adonis

| pairs | SumOfSqs | F.Model | R2 | p.value | p.adjusted |
|---|---|---|---|---|---|
| Syn3m vs. SoilCom+Syn3m | 0.821285 | 8.459117 | 0.18207658 | 0.001 | 0.01 |
| Syn3m vs. Syn2w | 5.981387 | 364.943267 | 0.90569392 | 0.001 | 0.01 |
| Syn3m vs. SoilCom+Syn2w | 5.09522 | 89.332686 | 0.70156916 | 0.001 | 0.01 |
| Syn3m vs. SoilCom | 5.986696 | 36.02102 | 0.48663231 | 0.001 | 0.01 |
| SoilCom+Syn3m vs. Syn2w | 5.169615 | 53.295959 | 0.58377128 | 0.001 | 0.01 |
| SoilCom+Syn3m vs. SoilCom+Syn2w | 4.245157 | 30.841391 | 0.44800651 | 0.001 | 0.01 |
| SoilCom+Syn3m vs. SoilCom | 4.060587 | 16.452385 | 0.3021426 | 0.001 | 0.01 |
| Syn2w vs. SoilCom+Syn2w | 0.15041 | 2.641276 | 0.06498998 | 0.096 | 0.096 |
| Syn2w vs. SoilCom | 6.40396 | 38.552621 | 0.50360942 | 0.001 | 0.01 |
| SoilCom+Syn2w vs. SoilCom | 5.463732 | 26.425971 | 0.41017575 | 0.001 | 0.01 |
